# Phylogenomic diversity elucidates mechanistic insights into Lyme borreliae host association

**DOI:** 10.1101/2022.05.25.493352

**Authors:** Matthew Combs, Ashley L. Marcinkiewicz, Alan P. Dupuis, April D. Davis, Patricia Lederman, Tristan A. Nowak, Jessica L. Stout, Klemen Strle, Volker Fingerle, Gabriele Margos, Alexander T. Ciota, Maria A. Diuk-Wasser, Sergios-Orestis Kolokotronis, Yi-Pin Lin

**Affiliations:** Department of Ecology, Evolution, and Environmental Biology, Columbia University, New York, NY USA; Division of Infectious Diseases, Wadsworth Center, NYSDOH, Albany, NY, USA; Department of Biomedical Sciences, SUNY Albany, Albany, NY, USA; German National Reference Centre for Borrelia at the Bavarian Health and Food Safety Authority, Oberschleissheim, Germany; Department of Epidemiology and Biostatistics, School of Public Health, SUNY Downstate Health Sciences University, Brooklyn, NY, USA; Institute for Genomics in Health, SUNY Downstate Health Sciences University, Brooklyn, NY, USA; Division of Infectious Diseases, Department of Medicine, College of Medicine, SUNY Downstate Health Sciences University, Brooklyn, NY, USA

**Keywords:** Host association, *Borrelia*, Complement, Phylogenomics, Plasmid diversity

## Abstract

Host association– the selective adaptation of pathogens to specific host species – evolves through constant interactions between host and pathogens, leaving a lot yet to be discovered on immunological mechanisms and genomic determinants. The causative agents of Lyme disease (LD) are spirochete bacteria composed of multiple species of the *Borrelia burgdorferi* sensu lato complex, including *B. burgdorferi* (*Bb*), the main LD pathogen in North America – a useful model for the study of mechanisms underlying host-pathogen association. Host adaptation requires pathogens’ ability to evade host immune responses, such as complement, the first-line innate immune defense mechanism. We tested the hypothesis that different host adapted phenotypes among *Bb* strains are linked to polymorphic loci that confer complement evasion traits in a host-specific manner. We first examined the survivability of 20 *Bb* strains in sera *in vitro* and/or bloodstream and tissues *in vivo* from rodent and avian LD models. Three groups of complement-dependent host association phenotypes emerged. We analyzed complement-evasion genes, identified *a priori* among all strains and sequenced and compared genomes for individual strains representing each phenotype. The evolutionary history of *ospC* loci is correlated with host-specific complement-evasion phenotypes, while comparative genomics suggests several gene families and loci are potentially involved in host association. This multidisciplinary work provides novel insights into the functional evolution of host adapted phenotypes, building a foundation for further investigation of the immunological and genomic determinants of host association.

**IMPORTANCE:** Host association is the phenotype that is commonly found in many pathogens that preferential survive in particular hosts. The Lyme disease (LD) causing agent, *B. burgdorferi* (*Bb*), is an ideal model to study host association, as *Bb* is mainly maintained in nature through rodent and avian hosts. A widespread yet untested concept posits that host association in *Bb* strains is linked to *Bb* functional genetic variation conferring evasion to complement, an innate defense mechanism in vertebrate sera. Here, we tested this concept by grouping 20 *Bb* strains into three complement-dependent host association phenotypes based on their survivability in sera and/or bloodstream and distal tissues in rodent and avian LD models. Phylogenomic analysis of these strains further correlated several gene families and loci, including *ospC*, with host-specific complement-evasion phenotypes. Such multifaceted studies thus pave the road to further identify the determinants of host association, providing mechanistic insights into host-pathogen interaction.

## INTRODUCTION

Infectious disease systems are governed by the evolution of host-pathogen interactions. Some pathogens can survive in multiple host species, but tend to preferentially adapt to some hosts over others – a process known as host association (Wolcott *et al*. 2021). Although this is an attractive theory, the mechanisms and genetic basis of host association remain largely unexplored. The genospecies complex of the bacterial spirochete *Borrelia burgdorferi* sensu lato (also known as *Borreliella burgdorferi* sensu lato or Lyme borreliae) are the causative agents of Lyme disease, the most common vector-borne disease in North America and Europe (Steere *et al*. 2016; Kugeler *et al*. 2021). Transmitted by a generalist *Ixodes* tick and carried by multiple vertebrate host species, the Lyme disease bacterium is a suitable model to study the mechanisms that mediate host association (Tufts *et al*. 2019; Wolcott *et al*. 2021). In fact, the field and laboratory evidence suggest different Lyme borreliae species vary in their host association (Brisson & Dykhuizen 2004; Steere *et al*. 2016; Tufts *et al*. 2019; Wolcott *et al*. 2021) For example, birds and rodents are the most common reservoir hosts that are selectively associated with *B. garinii* and *B. afzelii*, respectively, two frequently observed Lyme borreliae species in Eurasia (Tufts *et al*. 2019; Wolcott *et al*. 2021). In contrast, both host types were found to carry *B. burgdorferi* sensu stricto (hereafter *B. burgdorferi*), the most commonly isolated Lyme borreliae species in North America (Tufts *et al*. 2019; Wolcott *et al*. 2021). However, *B. burgdorferi* exhibits extensive strain diversity with different genotypes defined by several polymorphic loci (e.g., RST, MLST, *ospC*) (Theisen *et al*. 1995; Wang *et al*. 1999; Margos *et al*. 2010; Schwartz *et al*. 2021), and rodent or bird host associations have been documented in some genotypes of this spirochete species (Brisson & Dykhuizen 2004; Mechai *et al*. 2016; Lin *et al*. 2022). Thus, though still under debate, *B. burgdorferi* strains with different genotypes appear to exhibit variable host association.

To invade a host, Lyme borreliae require the ability to initially colonize and replicate at the tick bite site, migrate from those sites to the bloodstream, and subsequently disseminate to distal tissues (Steere *et al*. 2016; Radolf *et al*. 2020). In humans, systemic infections cause multiple manifestations in heart, joints, and neurological tissues, but in reservoir animals, the spirochetes persist at those distal tissues and organs without triggering symptoms (Tracy & Baumgarth 2017; Radolf *et al*. 2020). Systemic spread requires the mechanisms that facilitate hematogenous dissemination. As the first line of the host immune mechanism present in the blood, complement has been shown to control the ability of Lyme borreliae to disseminate to distal tissues (Kurtenbach *et al*. 2002; Dulipati *et al*. 2020; Lin *et al*. 2020a), emphasizing the role of this immune response in potentially dictating host association of Lyme borreliae.

Complement is a cascade comprised of several serum proteins, which can be activated by three canonical pathways (i.e. classical, alternative, and lectin pathways), resulting in digestion of these proteins to form different protein complexes (Reis *et al*. 2019). The activation of complement leads to the release of complement peptides, resulting in inflammation and phagocytosis. That activation also causes the deposition of several complement proteins (C5b, C6, C7, C8, and C9) that generate a pore-forming membrane attack complex (C5b-9) on the pathogen surface to lyse pathogens (Reis *et al*. 2019). In the absence of pathogens, vertebrate hosts produce complement regulatory proteins, such as factor H (FH), inhibiting the complement to prevent host cell damages (Zipfel & Skerka 2009; Gialeli *et al*. 2018). Similar to other pathogens, Lyme borreliae equip their surface with a group of anti-complement proteins to survive in the serum of the blood, known as serum resistance/serum survivability (Dulipati *et al*. 2020; Lin *et al*. 2020a; Skare & Garcia 2020). These proteins include CspA (a member of the protein family 54, Pfam54), CspZ, and OspE paralogs (collectively known as <u>c</u>omplement regulator <u>a</u>cquiring <u>s</u>urface <u>p</u>roteins, CRASPs) that bind to FH to inactivate the alternative complement pathway (Lin *et al*. 2020b). Additionally, the spirochete proteins BBK32 and Elp paralogs bind to C1 whereas the other spirochete protein, OspC, binds to C4b. These proteins block the activation of classical complement pathway whereas OspC also inactivates lectin complement pathway by binding to respective complement components (Garcia *et al*. 2016; Xie *et al*. 2019; Lin *et al*. 2020c; Pereira *et al*. 2021). Therefore, this variety of spirochete surface-localized, anti-complement proteins suggests these proteins, if polymorphic, confer Lyme borreliae species-or strain-specific host association. In fact, CspA is highly polymorphic among the variants from different Lyme borreliae species, and the allelic variation of this protein promotes rodent- or bird-specific complement evasion and spirochete transmission via tick feeding (Hart *et al*. 2021).

While divergent across Lyme borreliae species, the CspA variants from different strains within the same species (e.g. *B. burgdorferi*, *B. afzelii*) are highly conserved (∼99% sequence identity) (Hart *et al*. 2021). However, comparative analysis of *B. burgdorferi* genomes revealed a variable plasmid content and extensive protein-coding polymorphism (Qiu *et al*. 2004; Caine & Coburn 2016; Hart *et al*. 2021). These findings suggest that polymorphisms and the presence or absence of Lyme borreliae proteins with critical contributions to infectivity (e.g., anti-complement proteins) may confer *B. burgdorferi* strain-specific host association. To address this, we investigated the following: 1) Do different *B. burgdorferi* strains exhibit variable host-specific complement evasion phenotypes, and do they correlate with phylogenetic divergence at anti-complement proteins? 2) If so, do host-specific complement evasion phenotypes among strains link to distinct host-specific infectivity (i.e., tissue dissemination)? 3) Does genomic variation among strains with host-specific complement evasion phenotypes provide additional insights into mechanisms of host association? In this study, we examined the ability of multiple genotypically distinct *B. burgdorferi* strains to evade the complement from laboratory rodent and avian model animals, *Mus musculus* mouse and *Coturnix* quail, respectively, and promote infectivity in these hosts. We contextualized the results with phylogenetic analysis of anti-complement genes and comparative genomic analyses from these *B. burgdorferi* strains to provide mechanistic insights into Lyme borreliae host association.

## RESULTS

### 1. Phylogenetic divergence of spirochete anti-complement proteins corresponds with *B. burgdorferi* host-dependent anti-complement phenotypes

We first examined whether genotypically distinct *B. burgdorferi* strains display different levels of complement evasion phenotypes in a host-dependent manner, and if such differences are associated with the phylogeny of the anti-complement proteins from each of these strains. We thus chose 20 *B. burgdorferi* strains with different genotypes based on typing different polymorphic loci (i.e. *ospC*, RST, and MLST)(Table 1).

**Table 1.**
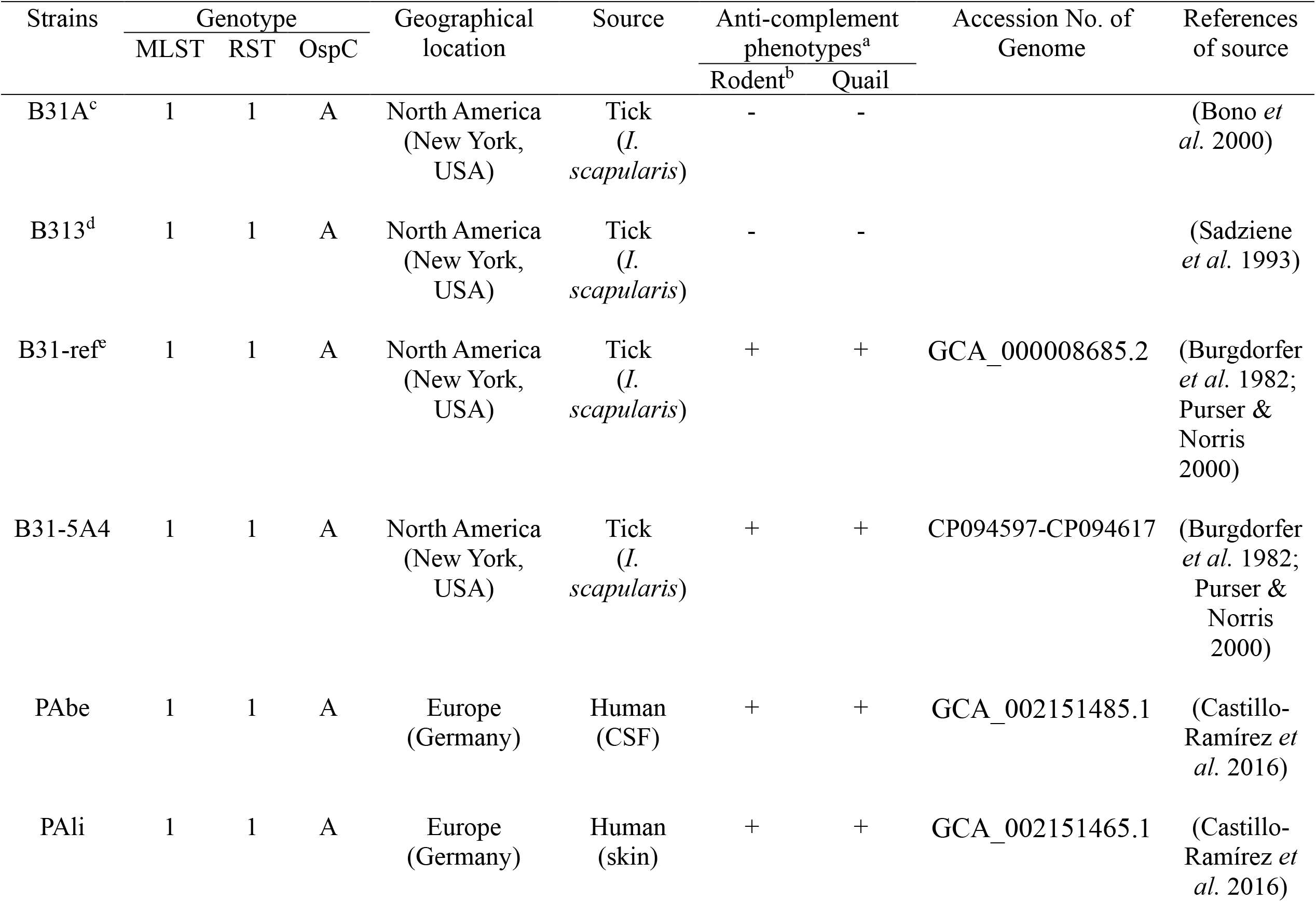
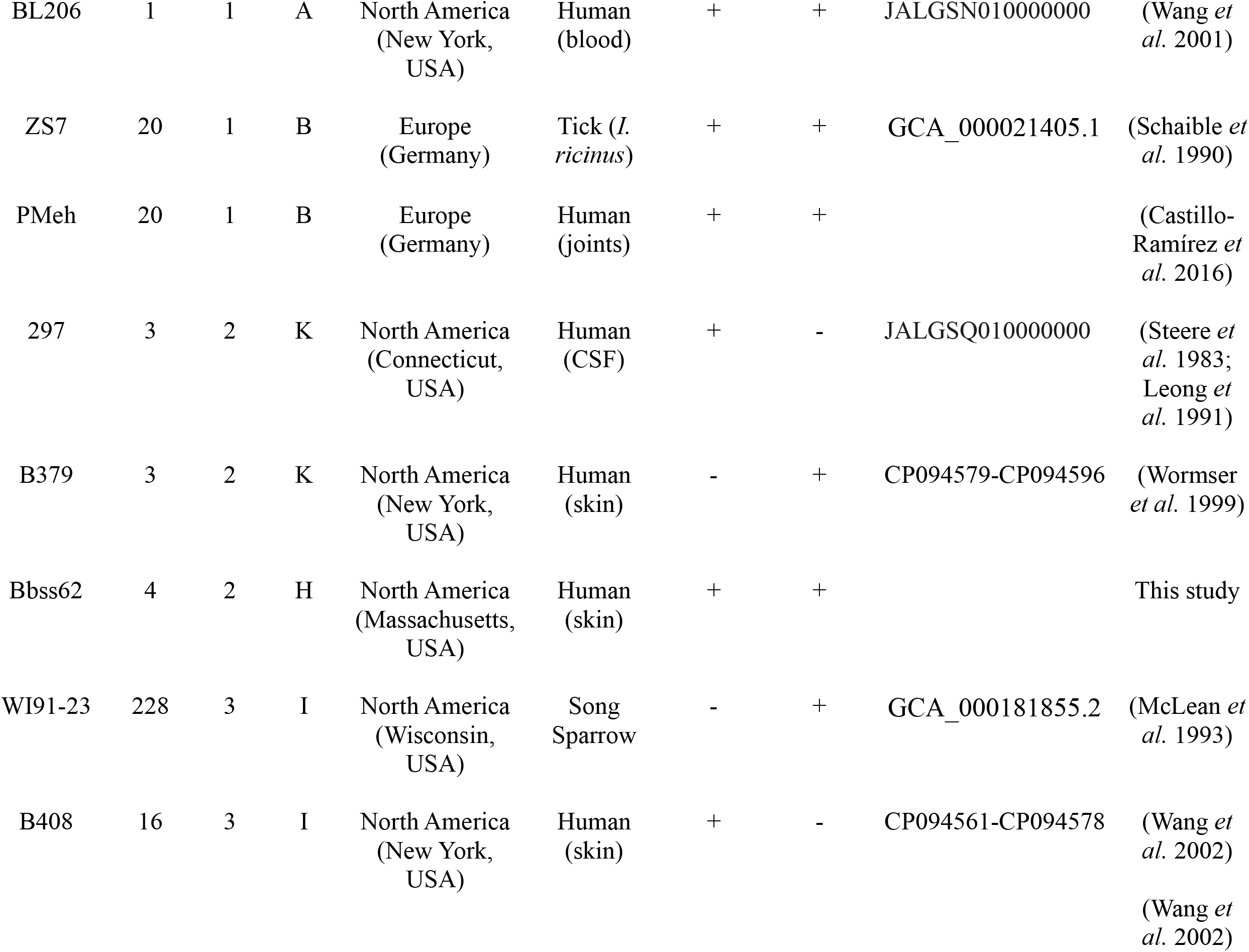
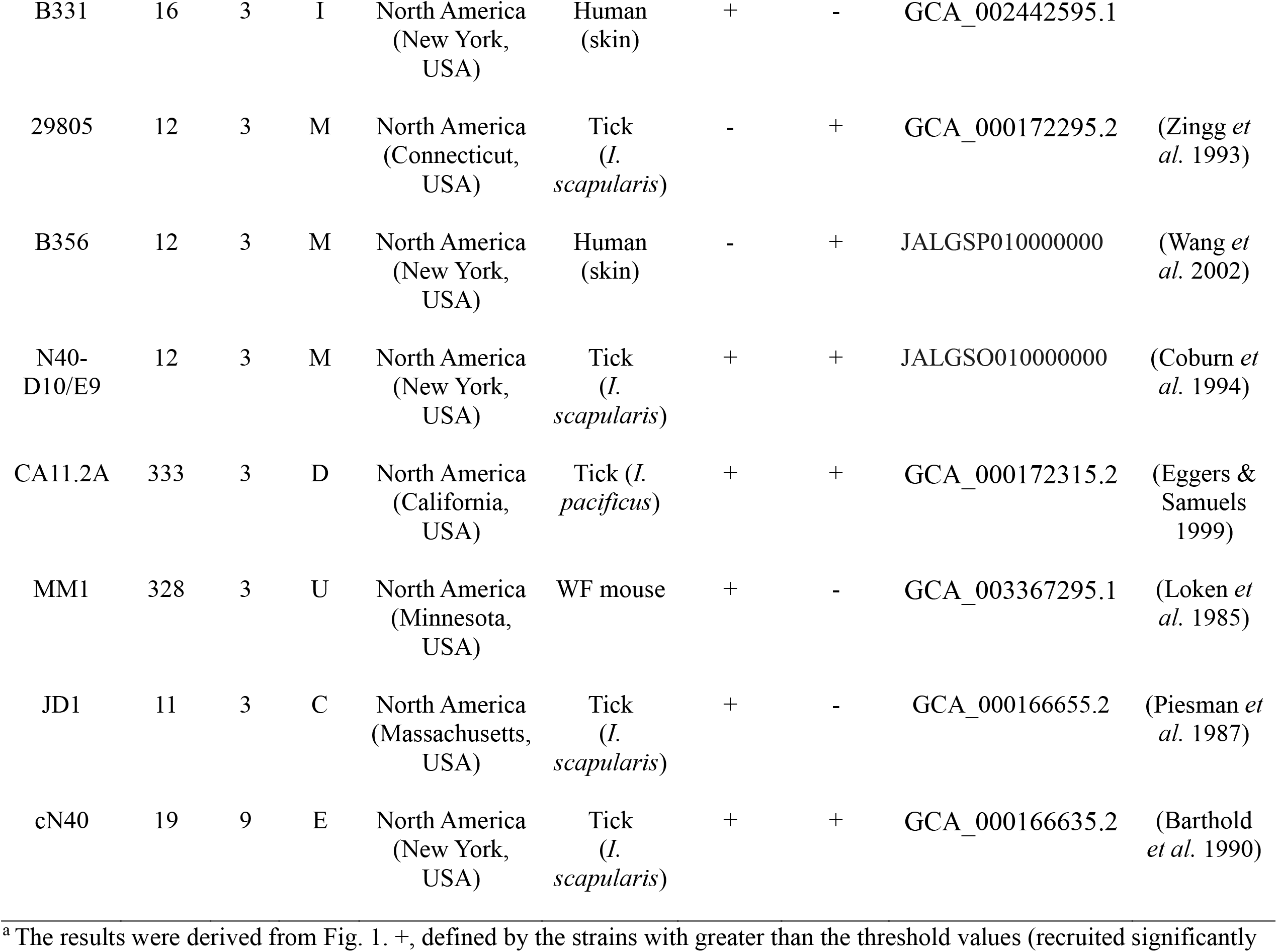
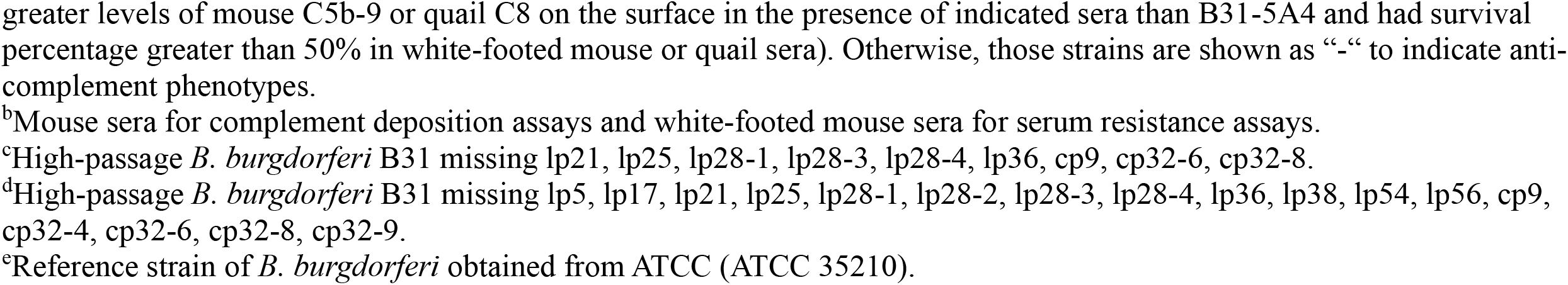
The genotypes and anti-complement phenotypes of *B. burgdorferi* strains used in this study.

#### 1.1 *B. burgdorferi* displays strain-level and host-specific variation in complement inactivation and serum resistance

We first added each of these strains, along with a serum-sensitive and complement-susceptible *B. burgdorferi* strain B313 (control) into the tested animal sera. These sera included the sera from BALB/c house mice (*Mus musculus*) and common quail (*Coturnix coturnix*), the rodent and avian models in the Lyme disease system, respectively (Hart *et al*. 2021). We then evaluated the surface deposition levels of mouse C5b-9 and quail C8 that are involved in bacterial lysis as the readout of complement activation. The deposition of mouse C5b-9 and quail C8 was apparent in B313, as expected (172 and 105 MFI for C5b-9 and quail C8 deposition, respectively)(Fig. 1A-D), in agreement with the inability of this strain to inactivate mouse and quail complement (Hart *et al*. 2018; Marcinkiewicz *et al*. 2019a). In contrast, the levels of the deposition for these complement proteins were close to undetectable on the surface of strain B31-5A4 (6 and 5 MFI for C5b-9 and quail C8 deposition, respectively), consistent with the fact that B31-5A4 inactivates both mouse and quail complement efficiently (Hart *et al*. 2018). We also found that the rest of the tested strains can be grouped according to their extent of complement deposition compared with that from B31-5A4 (Fig. 1A-D): <u>(a)</u> The strains displayed indistinguishable levels of deposition for both mouse C5b-9 and quail C8, suggesting their versatile ability to evade complement from both hosts (B31ref, PAbe, PAli, BL206, ZS7, PMeh, BBss62, N40-D10/E9, CA11.2A, and cN40), similar to B31-5A4 (Fig. 1C-D, red bars); <u>(b)</u> The strains exhibited significantly higher levels of mouse C5b-9 but not quail C8, suggesting the ability of the quail-specific complement evasion by these strains (B379, WI91-23, 29805, and B356) 5A4 (Fig. 1C-D, green bars). <u>(c)</u> The strains displayed significantly higher levels of quail C8 but not mouse C5b-9, suggesting mouse-specific complement evasion by these strains (297, B408, B331, MM1, and JD1) (Fig. 1C-D, blue bars).

We then examined the ability of these strains to survive in complement-containing sera of white-footed mouse (*Peromyscus leucopus*), the natural reservoir host of *B. burgdorferi* in North America, and quail sera. Note that the sera from white-footed rather than BALB/c mice were used to represent rodent sera because the complement in *Mus musculus* sera (i.e., BALB/c mouse sera) is labile *in vitro* (Caine & Coburn 2015; Ristow *et al*. 2015). As expected, 3 and 12% of B313 survived in white-footed mice and quail sera, respectively, consistent with this strain as serum-sensitive to both sera reported in recent studies (Fig. 1E-J) (Marcinkiewicz *et al*. 2019a; Lin *et al*. 2022). The survivability of all tested strains varies, ranging from 1.7% to 99.01%. To compare the strain’s ability in surviving in white-footed mouse or quail sera, we set up an arbitrary threshold of survivability at the middle (50%) (black dotted lines in Fig. 1E-J). Such a threshold allows us to group those strains into <u>(a)</u> the strains that survived in both white-footed mouse and quail sera at levels ≥50% (B31ref, B31-5A4, PAbe, PAli, BL206, ZS7, PMeh, BBss62, N40-D10/E9, CA11.2A, and cN40)(Fig. 1E, H, red bars); <u>(b)</u> the strains that survived in quail but not in white-footed mouse sera at levels ≥50% (B379, WI91-23, 29805, and B356) (Fig. 1E, H, green bars); <u>(c)</u> the strains that survived in white-footed mouse but not in quail sera at levels ≥50% (297, B408, B331, MM1, and JD1) (Fig. 1E, 1H, blue bars). All tested strains survived in complement-inactivated sera (white-footed mouse sera treated with cobra venom factor (CVF) and quail sera treated with *Ornithodorus moubata* complement inhibitor (OmCI)) at the levels greater than 90% (Fig. 1F-G, 1I-J). These results of grouping by serum survivability match the grouping by the levels of complement deposition showing three different complement evasion phenotypes for any tested strains (Table 1). Overall, these findings indicate a strain-specific ability to survive in white-footed mouse or quail sera and to prevent complement deposition, grouping genotypically distinct Lyme borreliae strains based on their host-specific anti-complement phenotypes.

**Figure 1.**
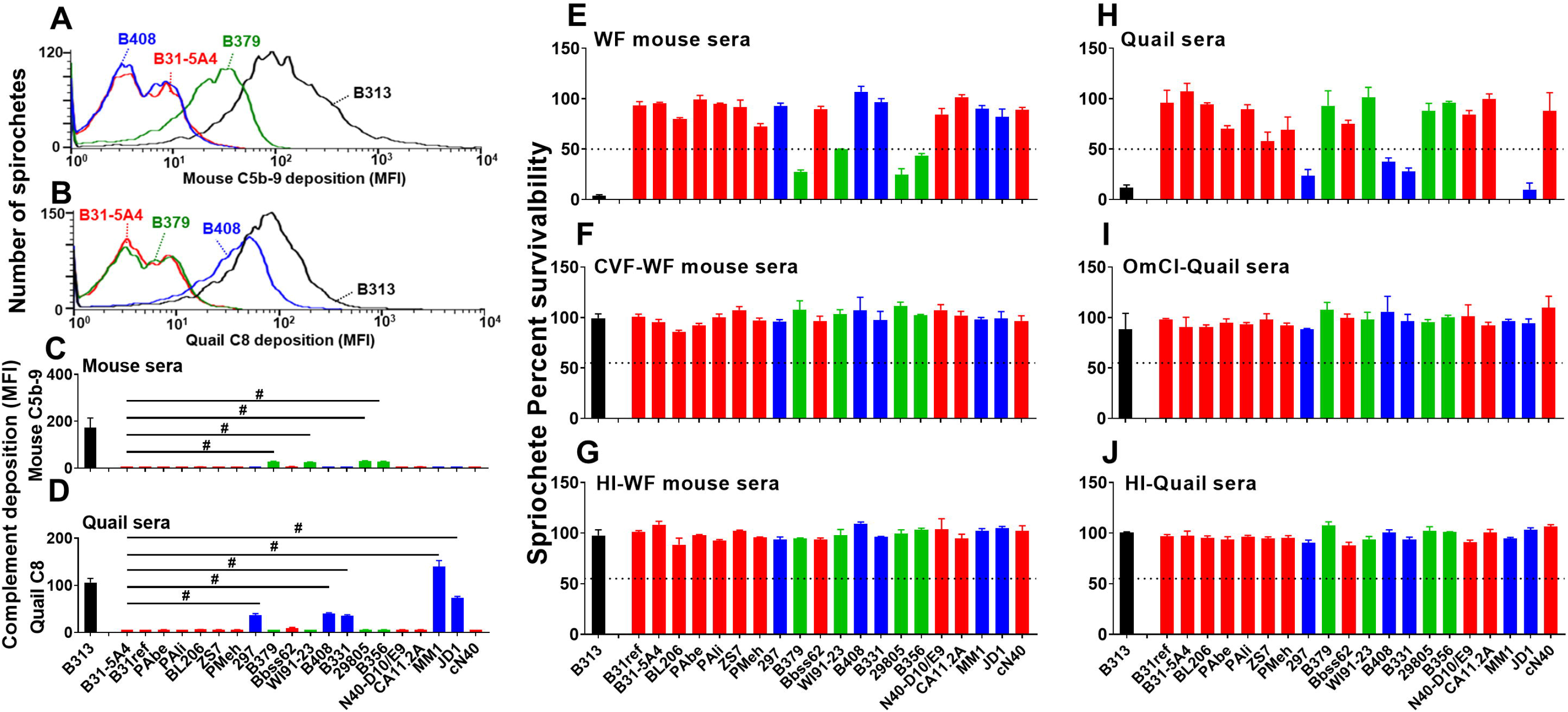
*B. burgdorferi* displays strain-to-strain variable ability of mammalian and avian serum resistance and complement inactivation. **(A to D)** Each of the indicated *B. burgdorferi* strains or the high passage, non-infectious, and serum sensitive *B. burgdorferi* strain B313 (control) was incubated with serum from mouse or quail with a final concentration of 20%. The bacteria were stained with the antibodies that recognize mouse C5b-9 or quail C8 prior to being applied to flow cytometry analysis as described in Materials and methods. Shown are the representative histograms of flow cytometry analysis presenting the deposition levels of **(A)** mouse C5b-9 or **(B)** quail C8 on the surface of indicated *B. burgdorferi* strains. The deposition levels of **(C)** mouse C5b-9 or **(D)** quail C8 on the surface of *B. burgdorferi* were measured by flow cytometry and presented as mean fluorescence index (MFI). Each bar represents the mean of three independent determinations ± SEM. Significant differences (p *<* 0.05, Kruskal-Wallis test with the two-stage step-up method of Benjamini, Krieger, and Yekutieli) in the deposition levels of mouse C5b-9 or quail C8 relative to the B31-5A4 were indicated (‘‘#’’). **(E to J)** Indicated *B. burgdorferi* strains were incubated for four hours with untreated sera from **(E)** white-footed (WF) mice or **(H)** quail, **(F)** CVF-treated white-footed mouse sera, **(I)** OmCI-treated qual sera, or heat inactivated sera from **(G)** white-footed mice or **(J)** quail. The number of motile spirochetes was assessed microscopically. The percentage of survival for those *B. burgdorferi* strains was calculated using the number of mobile spirochetes at four hours post incubation normalized to that prior to the incubation with serum. Each bar represents the mean of three independent determinations ± SEM. The black dotted lines indicate the threshold of percent survivability (50%). Bars are color coded to represent strains that can **(C to D)** efficiently inactivate complement from mouse (blue), quail (green), or both hosts (red) or **(E to J)** result in more than 50% survivability in the sera from white-footed mice (blue), quail (green) or both hosts (red).

#### 1.2 Phylogeny correlates *ospC* variation with the generalist vs. specialist anti-complement phenotypes of *B. burgdorferi*

We next investigated the phylogenetic relationships of the loci encoding anti-complement proteins from the above-mentioned *B. burgdorferi* strains. We reconstructed the phylogenetic history of the sequences of the documented anti-complement genes (*cspA*, *bbk32*, *ospC*, and *cspZ*) and the set of core chromosomal genes shared among all strains (control) from each of the tested *B. burgdorferi* strains. The phylogenetic correlation with the anti-complement phenotypes of these strains was evaluated. Although the proteins encoded by *ospE* and *elp* paralogs also display anti-complement phenotypes (Hellwage *et al*. 2001; Metts *et al*. 2003; Kraiczy *et al*. 2004; Pereira *et al*. 2021), we did not include them because they are located on the highly homogenous family of 32kb circular plasmids (cp32), which cause low-confidence assemblies derived from short-read sequences (Casjens *et al*. 1997a). All the anti-complement and core chromosomal gene phylogenies exhibited paraphyletic groups of strains with divergent anti-complement activity phenotypes (Fig. 2). Across all trees, terminal groups often contained both mouse-and quail-specific anti-complement phenotypes (Fig. 2, strains with mouse- and quail-specific anti-complement phenotypes are presented in blue and green, respectively; strains with versatile anti-complement phenotypes are shown in red). The phylogenies for *cspA*, *bbk32*, and *cspZ*, produced inconsistent correlations with Pagel’s λ distributions that ranged from zero to one (Fig. 2A-C, Fig. S1A-C). The *ospC* phylogeny exhibited a consistent, strong signal of non-random trait evolution where the anti-complement activity phenotype correlated was associated with tip position across the tree (λ=0.88), suggesting an association between *ospC* genotype and host-specific anti-complement activity (Fig. 2D, Fig. S1D). The phylogeny produced from the core set of chromosomal genes also exhibited a consistent signal of correlation with anti-complement activity phenotype (λ=0.93), suggesting that this phenotype may be driven by genome-wide evolution.

**Figure 2.**
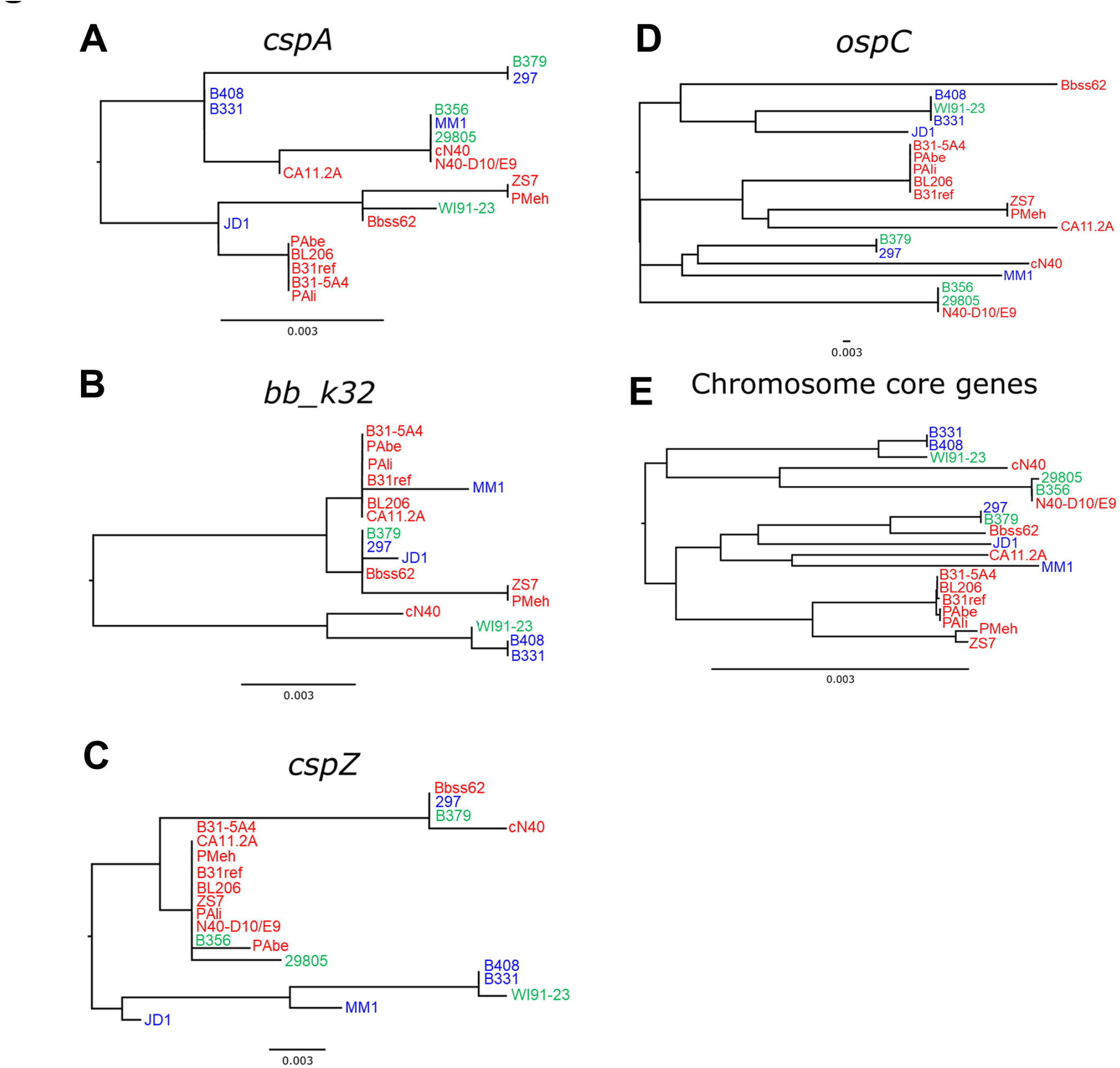
Phylogenetic trees of *B. burgdorferi* anti-complement proteins associate *ospC* and chromosomal core genes with hosts-specific complement inactivation activity. Individual phylogenies of **(A)** *cspA*, **(B)** *bb_k32*, **(C)** *cspZ*, **(D)** *ospC*, and **(E)** chromosome core genes represent evolutionary relationships among 20 indicated *B. burgdorferi* strains with known complement evasion phenotypes. Labels are color coded to represent strains that can efficiently inactivate complement from mouse (blue), quail (green), or both hosts (red). Strains that do not harbor particular loci are excluded from those respective trees.

### 2. Strain-variable, host-specific anti-complement phenotypes are linked to distinct levels of host-specific dissemination and genomic variation

Our findings associated the host-specific, anti-complement phenotypes with the phylogeny of one anti-complement protein from various genotypes of *B. burgdorferi* strains. These results raise several intriguing questions: 1) Do genotypically distinct *B. burgdorferi* strains confer complement-dependent infectivity that varies among host types? 2) If so, is genomic divergence associated with such an immunological variability of the phenotype *B. burgdorferi*? To address these questions, we chose a strain from the group displaying either quail- or mouse-specific anti-complement phenotypes (B379 and B408, respectively) along with the strain with a versatile ability to inactivate complement from both host types (B31-5A4).

#### 2.1 Genotypically distinct *B. burgdorferi* strains promote host-specific and complement-dependent phenotypes *in vivo*

##### 2.1.1 Strain-specific complement evasion activity promotes a host-specific variation in *B. burgdorferi* bloodstream survivability

The presence of complement in host bloodstream raises the hypothesis of strain-specific anti-complement phenotypes conferring host-dependent *B. burgdorferi* bloodstream survivability. We thus examined the ability of strains B31-5A4, B379, and B408 to survive in the mouse or quail bloodstream 1-h post intravenous (iv.) injection (hpi), a model established to test the short-term ability of *B. burgdorferi* to trigger bacteremia (Caine & Coburn 2015). In the blood from BALB/c mice, strains B31-5A4 and B408 yielded significantly greater levels of bacterial burdens than B379 and the negative control strain, B31A (a high-passage strain that cannot efficiently survive in vertebrate bloodstream (Caine & Coburn 2015)) (Fig. 3A). Conversely, in quail blood, B31-5A4 and B379 induced significantly higher levels of bacterial loads than B408 and the control strain B31A (Fig. 3B). We observed indistinguishable burdens of these strains in the blood from C3^-/-^ BALB/c mice or OmCI-injected quail at 1hpi (Fig. 3C-D). These results correlated the bloodstream survivability of *B. burgdorferi* strains with these strains’ anti-complement phenotypes in a host-specific manner.

**Figure 3.**
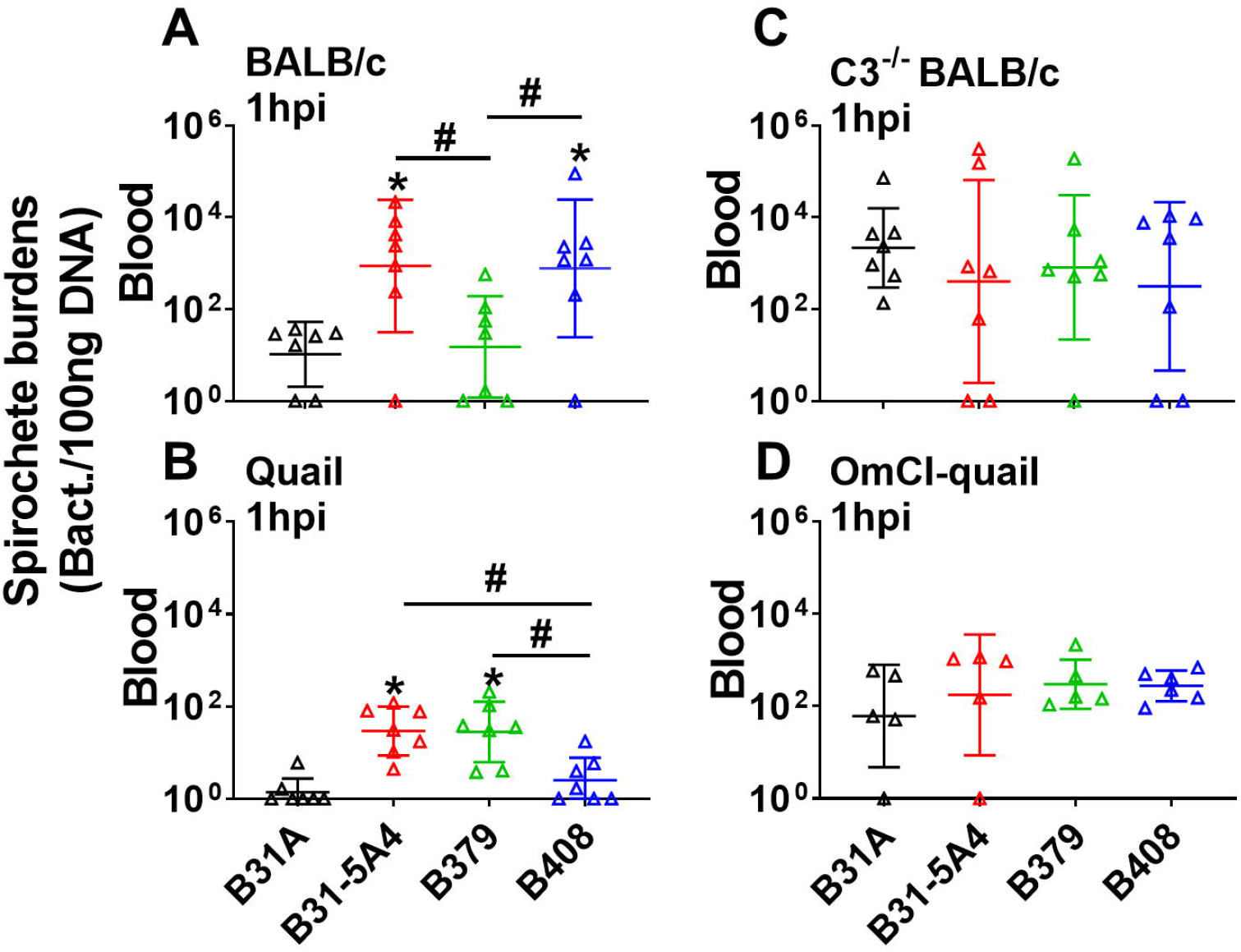
Complement dictates mouse and quail-specific short-term bloodstream survival of *B. burgdorferi* in a genotype-dependent manner. **(A)** BALB/c mice, **(B)** untreated quail, **(C)** C3^-/-^ mice in BALB/c background, or **(D)** OmCI-treated quail were iv. inoculated with *B. burgdorferi* strains B31-5A4, B379, or B408, or a high passage, non-infectious *B. burgdorferi* strain B31A (control) (Five animals per group for OmCI-treated quail and seven animals per groups for others). Blood was collected from these animals at 1-hr after inoculation, and bacterial burdens were quantified by qPCR. Shown are the geometric mean of bacterial loads ± geometric standard deviation of five mice or quail per group. Significant differences (p < 0.05, the Kruskal-Wallis test followed by the two-stage step-up method of Benjamini, Krieger and Yekutieli.) in the spirochete burdens from the burdens in B31-infected animal blood (“*”), or between two strains relative to each other (‘‘#’’).

##### 2.1.2 Complement is essential to determine the host-dependent, *B. burgdorferi* strain-specific early onsets of dissemination

The ability of bacterial survival in the blood has been associated with phenotypes of disseminated infection (Caine & Coburn 2015; Lin *et al*. 2020c), we thus investigated this by generating *I. scapularis* nymphs carrying indistinguishable levels of B31-5A4, B379, or B408 (Fig. S2A). After allowing these nymphs to feed on BALB/c mice, we evaluated bacterial burdens in replete nymphs and the tissues at 10 days post tick feeding (dpf), the time point when spirochetes begin their systemic spread in mice, representing early onsets of dissemination (Hart *et al*. 2021) (Marcinkiewicz et al. unpublished data). We also measured bacterial burdens in the same fashion at 14dpf, the later time point of dissemination to distal tissues. After tick feeding, we found similar levels of these strains in replete nymphs (Fig. S2B). At 10dpf, B31-5A4, B379, and B408 displayed similar levels at the tick bite sites on skin (Fig. 4A), but the bacterial burdens of B379 were significantly lower than those of B31-5A4 or B408 in blood and all distal tissues (tibiotarsus joints, bladder, and heart; Fig. 4B-E). At 14dpf, we found significantly lower bacterial burdens in all tissues, including the tick bite sites on skin of B379-infected mice, compared to those from B31-5A4- or B408-infected mice (Fig. S3A-E). We also tested the role of complement in determining such a strain-specific dissemination by infecting the complement-deficient BALB/c mice (C3^-/-^ mice) in the same manner. Though bacterial burdens in all tissues from B379-infected mice remain significantly lower than those from B31-5A4- or B408-infected mice at 14dpf (Fig. S3F-J), the infection of all tested strains yielded similar levels of bacterial loads in all tissues and replete nymphs at 10dpf (Fig. 4F-J, S2D). These results indicate an essential role for complement in determining strain specificity at the beginning of dissemination.

**Figure 4.**
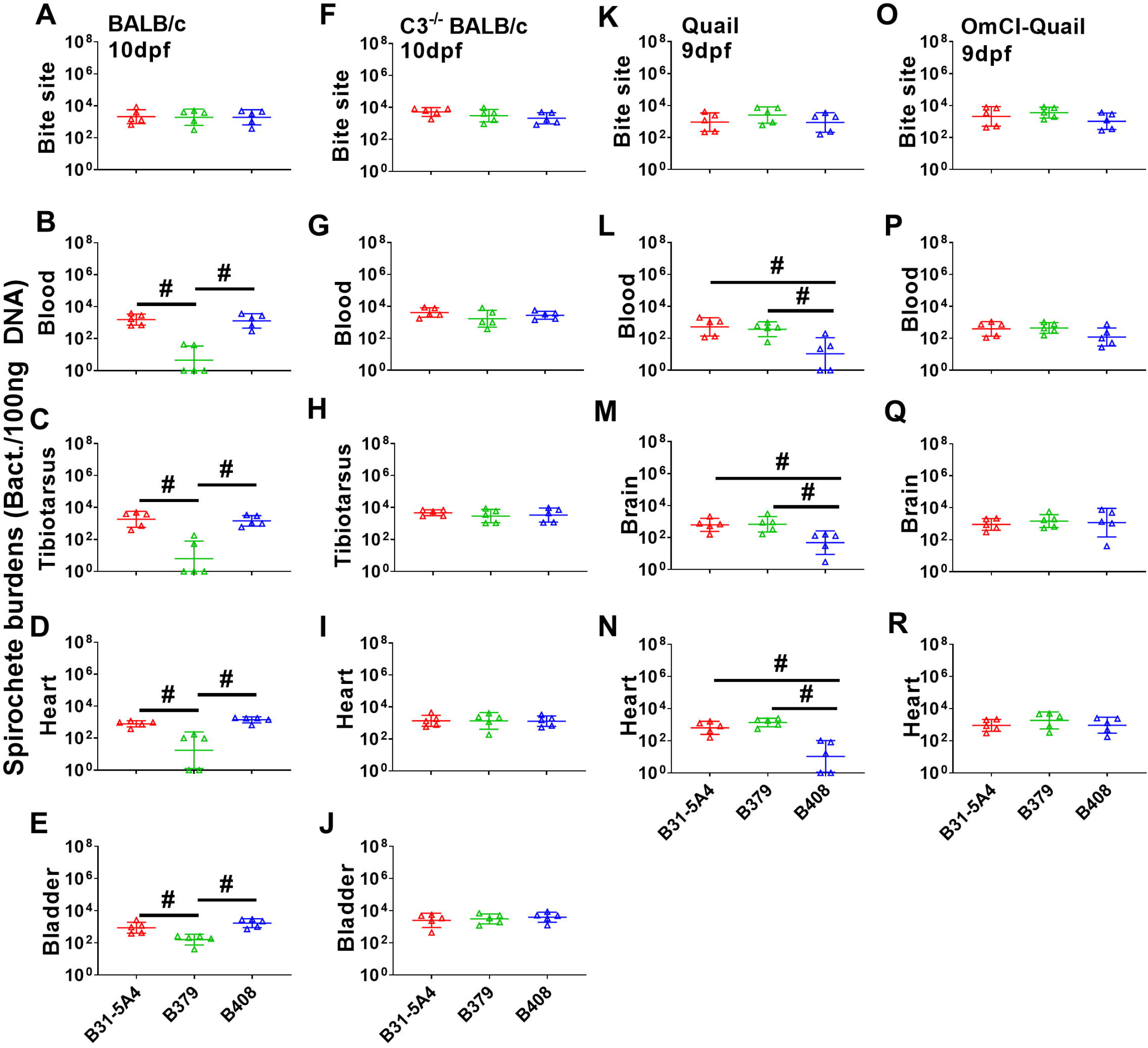
*B. burgdorferi* exhibits host- and bacterial genotype-specific early dissemination in a complement-dependent fashion. The *I. scapularis* nymphs carrying *B. burgdorferi* strains B31-5A4, B379, or B408, were allowed to feed until they are replete on five **(A to E)** wild-type or C3^-/-^ **(F to J)** BALB/c mice, **(K to N)** wild type or **(O to R)** OmCI-treated quail. The mice and quail were euthanized at 10- and 9-days post nymph feeding (dpf), respectively. The bacterial loads at **(A, F)** the site where nymphs fed (“inoc. site”), **(B, G)** blood, **(C, H)** tibiotarsus joints, **(D, I)** heart, and **(E, J)** bladder of mice and **(K, O)** the site of nymphs bite (“inoc. site”), **(L, P)** blood, **(M, Q)** brain, and **(N, R)** heart of quail collected immediately after euthanasia were determined by qPCR. The bacterial loads in tissues or blood were normalized to 100 ng total DNA. Shown are the geometric mean of bacterial loads ± geometric standard deviation of five mice or quail per group. Significant differences (p < 0.05, the Kruskal-Wallis test followed by the two-stage step-up method of Benjamini, Krieger and Yekutieli.) in the spirochete burdens between two strains relative to each other (‘‘#’’).

We then tested whether such a strain-specific, complement-dependent infectious phenotype at the beginning of dissemination is dependent on the host used in the study. We allowed the nymphs carrying B31-5A4, B379, or B408 to feed on quail and measured bacterial burdens at 9dpf, the time point when dissemination begins in quail, which represents the early infection onset for quail infected via tick feeding (Hart *et al*. 2021)(Marcinkiewicz et al. unpublished data). Quail-associated replete nymphs had similar levels of all tested strains, like mouse-associated replete nymphs (Fig. S2C). At 9dpf, all strains exhibited indistinguishable levels of bacterial burdens at the tick bite sites on skin (Fig. 4K), similar to the infection in mice. However, unlike in mice, the B408 infection resulted in lower bacterial loads in quail blood, and distal tissues (brain and heart), compared to B31-5A4 and B379 (Fig. 4L-N). Once those infected ticks fed on complement-deficient quail (OmCI-injected quail) for 9 days, we found indistinguishable levels of bacterial burdens in replete nymphs and any tested tissues (Fig. 4O-R, S2E). Together with the results from mice, these findings suggest an essential role of complement in dictating strain-specific, host-dependent early onsets of dissemination.

#### 2.2 Genomic divergence in *B. burgdorferi* strains with distinct host-specific anti-complement phenotypes

The host- and strain-specific, complement-dependent infectivity of B31-5A4, B379, and B408 raises the likelihood of differentiation in genome content, organization, and patterns of epigenetic modifications in these strains. We generated high-quality genome sequences and assemblies of strains B31-5A4, B379, and B408 using a long-read approach (HiFi reads with circular consensus sequencing on the Pacific Biosciences platform) (Fig. 5). While each genome contained a single highly homologous linear chromosome, they each exhibited a unique suite and number of plasmids. B31-5A4 contains 11 linear and 9 circular plasmids. B379 contains 8 linear and 9 circular plasmids, though an additional plasmid (lp21) appears fused to the 3’ end of its linear chromosome. B408 contains 10 linear and 7 circular plasmids. Gene annotations revealed a variable gene content, with 1527, 1435, and 1408 annotations identified in B31-5A4, B379, and B408 respectively (Table S1). We also identified methylation motifs, including two previously unreported motifs (GNAAGC in B379, GAAGG in B408). The methylation rates of each strain vary with B408 having the lowest rate among three strains (1.03 base methylations per 1000bp) (Table S2).

**Figure 5.**
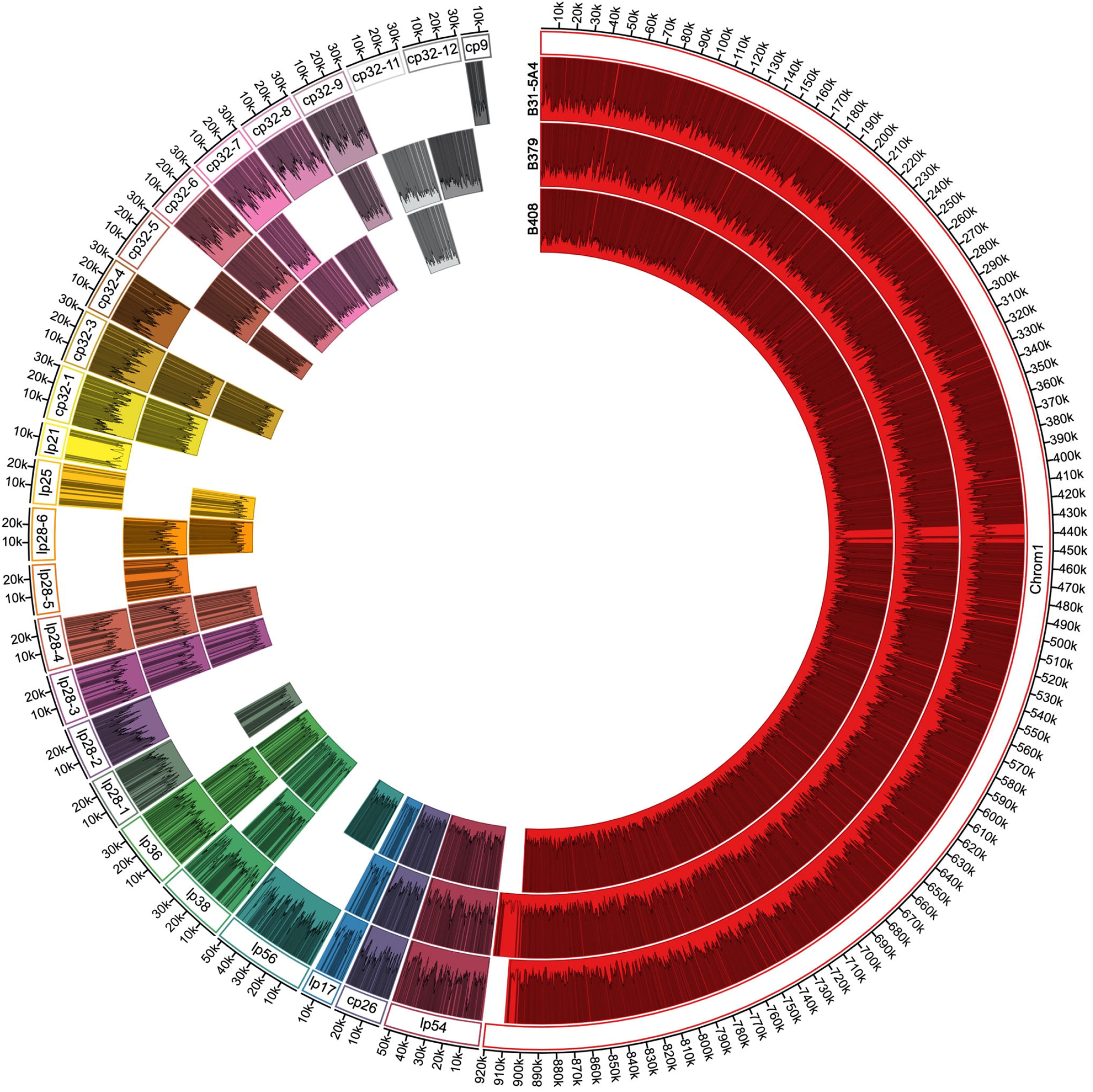
Genomic content of three *B. burgdorferi* strains. The genome of B. burgdorferi strains B31-5A4, B379, or B408 is represented by a concentric circle and each color-coded individual segment represents a genomic plasmid or chromosome, defined by labels provided in the outermost ring. Gaps in the ring represent plasmids that are not present in the genome at the time of sequencing. Grey lines within each segment represent gene annotations and the plots within each segment represent the number of methylated nucleotides per 1000 bases.

##### 2.2.1 Strain-specific duplication and deletion of loci including anti-complement genes

Many regions of high similarity between B31-5A4, B379, and B408 emerged, including all across the linear chromosome, circular plasmid 26 (cp26) and much of lp28-3, lp28-4, and lp54 (Fig. 5, 6A-C, S4). In contrast, we identified highly divergent gene content and organization along lp36. Both B379 and B408 exhibited a ca. 14kb deletion at the 3’ end of lp36, when compared with B31-5A4 (Fig. 6D). This deletion has removed an entire group of Pfam75 immunogenetic proteins, including P37 (i.e BB_K50), which are not found elsewhere in the genome (Fig. 6D). Similarly, B379 lp38 retained only 12 of the 35 genes found on lp38 of B31-5A4 and B408. The latter two strains appear to have a highly conserved lp38 (Fig. 6E). Additionally, the 5’ end of lp17 exhibits high variability; in B379 this plasmid exhibits a ca. 4kb fragment from lp36, whereas in B408 it contains a 2.6kb fragment from lp28-3 (Fig. 6F). Among other non-cp32 plasmids, few were found in a single strain (lp28-2 and cp9 only in B31-5A4; lp28-5 only in B379), while others were recovered from only two strains (lp56, lp28-1, and lp25 only absence in B379; lp28-6 only absence in B31-5A4) (Fig. 5).

**Figure 6.**
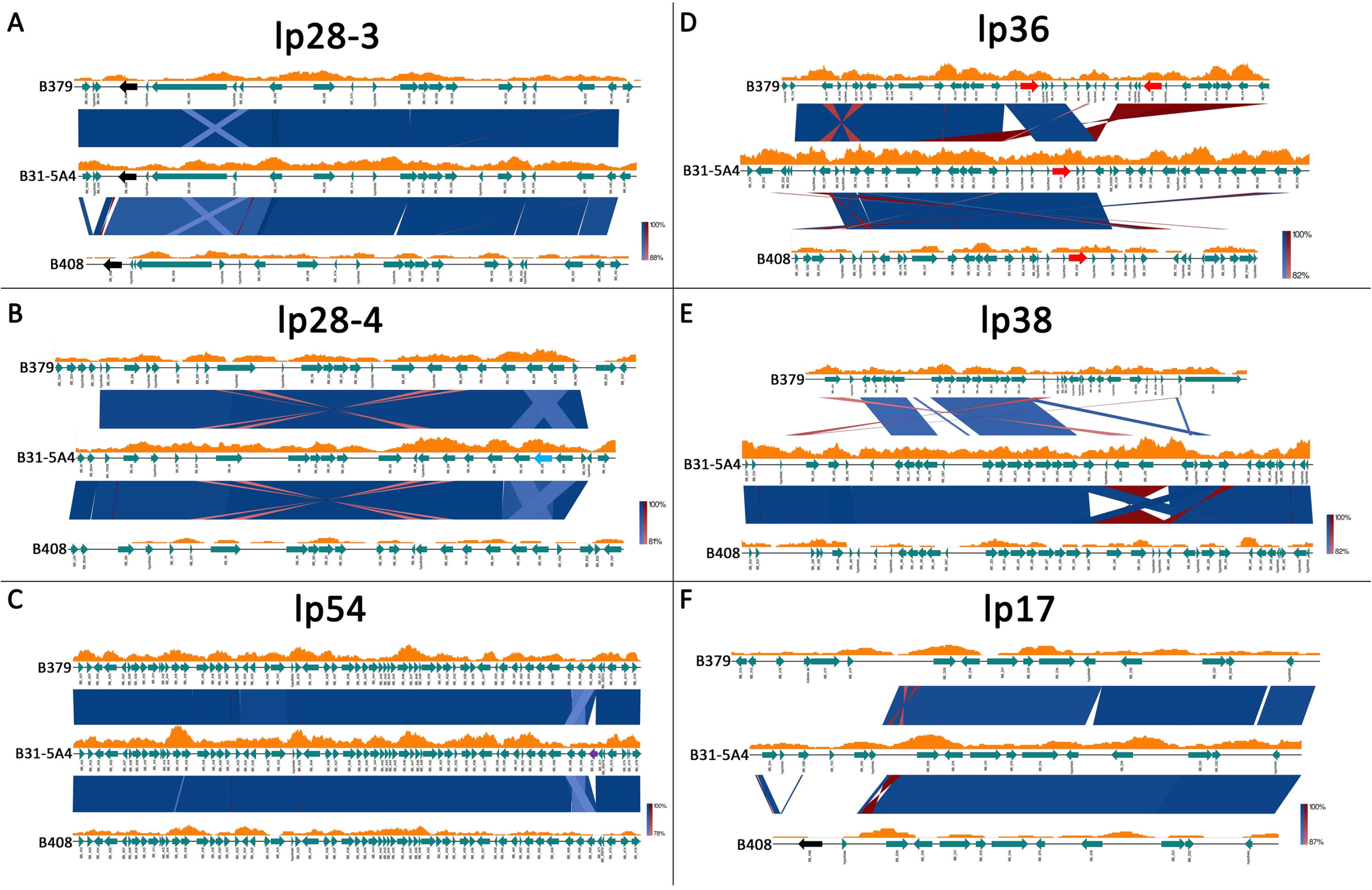
Genomic comparison reveals plasmid-specific differences among *B. burgdorferi* strains with distinct host-specific anti-complement phenotypes. The sequences of **(A)** lp28-3, **(B)** lp28-4, **(C)** lp54, **(D)** lp36, **(E)** lp38 and **(F)** lp17 from *B. burgdorferi* strains B31-5A4, B379, and B408 are represented by a black line and gene annotations are depicted by teal arrows, labeled according to homology to known genes in the reference genome of *B. burgdorferi* strain B31. Segments connecting each strain represent filtered BLAST results in either the same orientation (blue) or opposite orientation (red), with darker shades representing closer matches. Orange graphs above each strain depict the sliding window calculation of methylated nucleotides per 1000 bases. The colored arrows indicated the loci of *cspZ* (black), *bb_i38* (light blue), *bb_a70* (purple), *bb_k32* (red).

We also investigated gene content of specific anti-complement loci (*ospC*, *bb_k32*, *cspZ*, and *cspA*) and their associated gene families in B31-5A4, B379, and B408. Each strain harbors a single copy of *ospC* on cp26 (red arrows in Fig. S4). While B31-5A4 and B408 have a single copy of *bb_k32* on lp36, B379 carries two copies on lp36 (red arrows in Fig. 6D). Additionally, whereas B31-5A4 and B379 each possesses one copy of *cspZ* on lp28-3, B408 contains two copies with a second *cspZ* locus found on the 5’ end of lp17 (black arrows in Fig. 6F). While we found *cspA* as a single gene copy in all three strains, we observed several deletions of other Pfam54 members. *bb_a70* was absent from B379 and B408, but found on lp54 in B31-5A4 (purple arrow in Fig. 6C). Further, *bb_i38* was found on lp28-4 in B31-5A4, though the neighboring Pfam54 loci of B379 and B408 were retained (light blue in Fig. 6B). Taken together, these results showcase regions of high plasmid similarity and variability, along with evidence of locus duplication and deletion events, including those specific to anti-complement activity, across *B. burgdorferi* strains with distinct host-specific anti-complement phenotypes.

Across the 17 additional strains for which we tested serum survival phenotypes, variation in genome assembly quality and completeness prevent direct comparisons of plasmid organization, but permit gene content analysis in association with strain-specific phenotypes. Our paired Roary and Scoary analysis identified only a single gene with known function, immunogenic P37, that was found in generalist strains but not within the genome of any host-adapted strains.

##### 2.2.2 Characterization of variable genes with anti-complement determinants

Most loci showed 90-100% similarity a high degree of conserved functions (Fig. 7). Few loci exhibited ≤90% similarity, such as the highly polymorphic *ospC* and *dbpA* (Table S3) (Wilske *et al*. 1993; Roberts *et al*. 1998) and several genes documented to facilitate spirochete survival *in vitro* and *in vivo* and/or reduce infectivity after immunization, such as *bb_k13* and *vraA* (*bb_i16*) (Table S3) (Labandeira-Rey *et al*. 2001; Aranjuez *et al*. 2021). Among the genes encoding proteins with documented anti-complement phenotypes, *cspA*, *cspZ*, and *bbk32* differ by relatively few polymorphisms, averaging 99.29%, 96.91%, and 97.24% similarity among all strain pairs, respectively, while the average similarity among *ospC* loci was 85.80% (Table S3).

**Figure 7.**
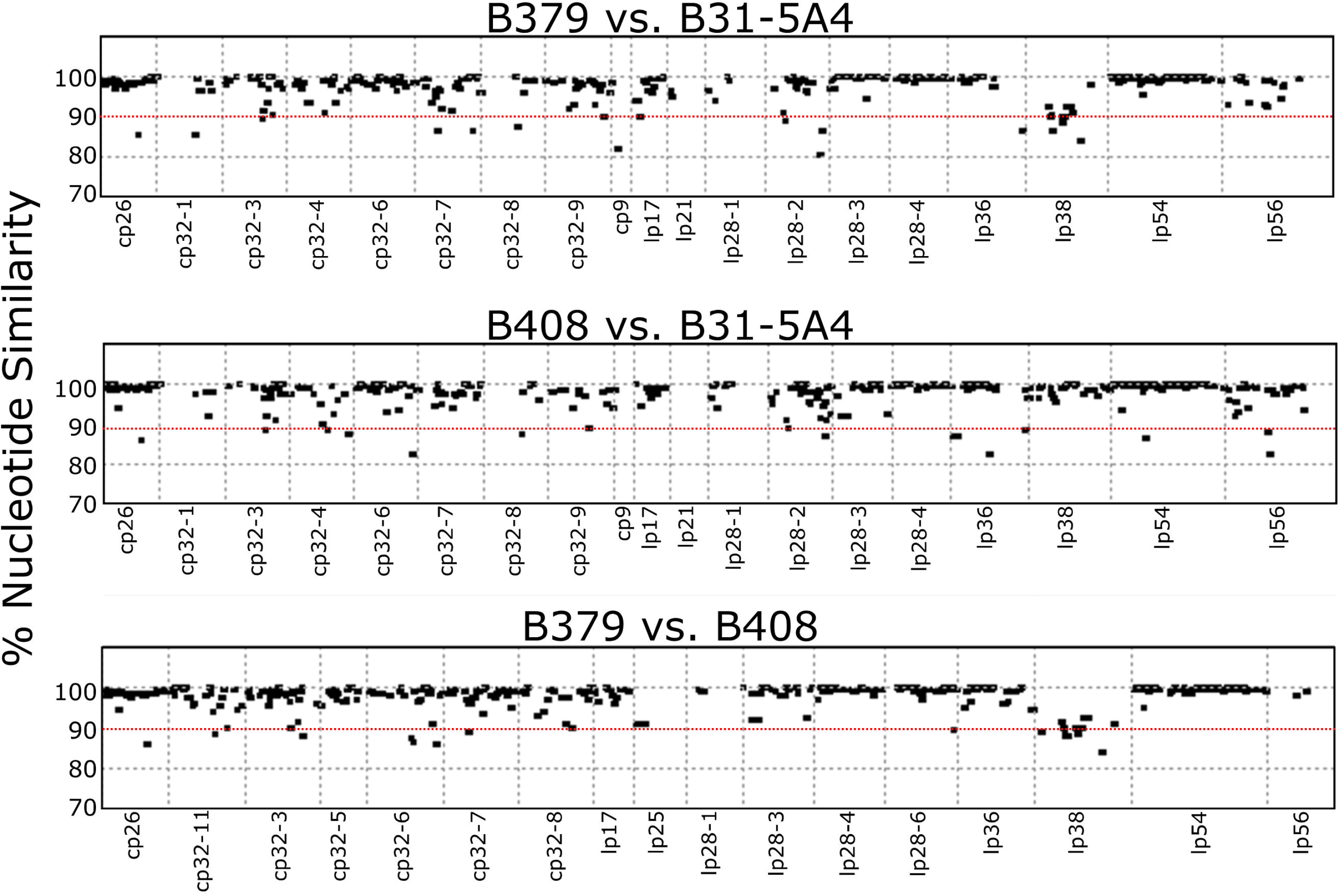
Loci-specific analysis identifies the polymorphic genes of the alleles among *B. burgdorferi* strains with distinct host-specific anti-complement phenotypes. Nucleotide similarity among annotated loci pairwise between **(A)** B379 and B31-5A4, **(B)** B408 and B31-5A4, and **(C)** B379 and B408. The x axis represents the position of each annotation within the latter of the two compared genomes. The y axis represents observed nucleotide similarity calculated using the nucmer program. The red dotted lines indicate the threshold (90% nucleotide similarity) to define the polymorphic loci.

## DISCUSSION

Differences in host association among pathogen species and strains are often the result of adaptive evolution by pathogens to host specific immune responses (Douam *et al*. 2015; Wolcott *et al*. 2021). The selective pressure imparted by host immune responses can result in the promotion of specific alleles or patterns of genomic variation in naturally occurring pathogens, known as multiple niche polymorphism (MNP) (Brisson & Dykhuizen 2004). Lyme borreliae exhibit strain-specific diversity in host range and genomic organization, with lipoprotein genes constituting a rapidly evolving part of the genome (Wywial *et al*. 2009; Qiu & Martin 2014; Tufts *et al*. 2019; Schwartz *et al*. 2021; Lin *et al*. 2022). Lipoproteins are associated with multiple cellular and immunological mechanisms influencing infection, thus making Lyme borreliae an ideal model for the investigation of the underlying mechanisms driving MNP-mediated host association. One such immunological mechanism is host-specific complement evasion that has been shown to differ among Lyme borreliae species, whereby particular species are adapted to evade host-specific complement (Kurtenbach *et al*. 1998, 2002; Lin *et al*. 2020a). In this study, our serum survivability and complement deposition assays demonstrated that *B. burgdorferi* strains exhibit either mouse-specific, quail-specific, or versatile complement evasion activity, extending the concept of host-specific complement evasion to different strains within one spirochete species.

Strains with quail-specific complement evasion disseminate more efficiently than other strains in wildtype quail during the early stages of dissemination (i.e., 9dpf), but are indistinguishable from others in complement-deficient quail. Similarly, more efficient dissemination phenotypes in wildtype mice were conferred by the strains with mouse-specific complement evasion, compared to other strains, during the beginning of mouse dissemination (i.e., 10dpf), but were indistinguishable from other strains in complement-deficient mice. These results are in agreement with the anti-complement activity documented to mediate early onset-specific dissemination of *B. burgdorferi* (Lawrenz *et al*. 2003; Woodman *et al*. 2007; Zhi *et al*. 2018). Therefore, our findings of strain-specific, host-dependent complement evasion activity and early-onset dissemination suggest that *B. burgdorferi* strains differ in their efficiency to disseminate, particularly at early stages, and such differences are host-specific and mediated by complement. Our findings also indicate that such a complement-dependent, strain-specific dissemination is less apparent at later time point of spirochete dissemination (i.e., 14dpf). In fact, additional mechanisms such as adaptive immune responses are involved in controlling spirochete colonization at later stages (Tracy & Baumgarth 2017; Bockenstedt *et al*. 2020). Such mechanisms may play more significant roles at later stages in diverse tissues, compared to complement. Thus, our results suggest the involvement of complement-independent mechanisms, impacting infection dynamics at tick bite sites on skin and distal tissues during later-onset (14dpf) stages of infection – warranting further investigation.

We characterized genetic variation related to these divergent phenotypes using multiple approaches. First, we examined phylogeny-phenotype correlations at targeted loci with known roles in complement evasion using publicly available and new short-read-based genome assemblies of variable completeness. We examined gene content and variability, plasmid organization, and epigenomic modifications in strains representative of each complement evasion phenotype using high accuracy long-read sequencing. Among the examined complement evasion loci, only *ospC* gene genealogy correlated with host-specific anti-complement phenotypes among tested spirochete strains. As associations between specific *ospC* types of *B. burgdorferi* strains and mammalian or avian reservoir animals have been observed (Brisson & Dykhuizen 2004; States *et al*. 2014; Vuong *et al*. 2014), our results raise a possibility that OspC is an anti-complement determinant that directly contributes to host-specific complement evasion and infectivity. That possibility is consistent with our serum resistance assays using naïve animal sera and OspC in binding C4b, interfering the impact of OspC-mediated evasion to an antibody-independent complement pathway (i.e., lectin pathway)(Caine *et al*. 2017). Additionally, this possibility is supported not only by the polymorphism of OspC among different *B. burgdorferi* strains but also by its complement ligand, C4b which varies among different host taxa (i.e. ∼35% sequence identity between mouse and quail) (Wilske *et al*. 1993). However, such a possibility is at odds with several findings that *B. burgdorferi* strains carrying identical *ospC* sequences display distinct infectivity in mice (e.g., the strains N40-D10/E9 vs. B356 (Wang *et al*. 2001; Chan *et al*. 2012) or B379 from this study and 297 (Masuzawa *et al*. 1992)). Thus, it is also possible that OspC serves as a marker, and other anti-complement loci linked to OspC in spirochetes contribute to such host-specific anti-complement phenotypes. Nonetheless, we did not find a correlation between host-specific complement evasion phenotypes and the phylogenies estimated for other tested anti-complement loci (i.e. *cspA*, *cspZ*, *bbk32*). These results suggest other anti-complement determinants (e.g., *ospE* and *elp* paralogs, not tested in this study) contribute to the complement evasion activity, or that each of these variants contribute to the host-specific complement evasion at different extents (i.e., polygenic effects) in different strain-host specific interactions. These possibilities are not mutually exclusive and would require future work to thoroughly examine.

The genomes of *B. burgdorferi* strains regularly recombine and gene repertoires vary, leading to unique plasmid profiles (Caine & Coburn 2016; Casjens *et al*. 2017). Gene duplication and deletion have been consistently found in *B. burgdorferi* genomes, which may influence adaptive traits (Kondrashov *et al*. 2002; Casjens *et al*. 2012). Consistent with these findings, we found evidence of recombination, duplication, and deletion in three representative *B. burgdorferi* strains with distinct host-specific phenotypes of hematogenous dissemination. These results suggest that the presence or absence of certain genes, or the differences in copy numbers for those genes caused by genetic modification events, impact distinct host-specific adaptation and eventually lead to *B. burgdorferi* diversification. For example, the members of Pfam75 are encoded on lp36 and expressed when spirochetes reside in hosts, though the roles of these members *in vivo* remain unclear (Fikrig *et al*. 1997; Hodzic *et al*. 2003). We found that the strains that efficiently disseminate in a single host in this study (i.e., B379 and B408) share a large 3’ end deletion of lp36, resulting in the absence of Pfam75 genes, whereas the strain that efficiently spreads in multiple hosts (i.e., B31-5A4) maintains intact Pfam75. This is consistent with the genome of *Borrelia garinii*, a species specifically adapted to bird, which lacks many lp36-encoded loci, including Pfam75 genes (Glöckner *et al*. 2004). Our findings indicate the need for future studies to examine role of these genes in determining host-specific infectivity.

When anti-complement genes were specifically examined in this comparative genomic study, we found that strains B379 and B408, with distinct host-specific dissemination phenotypes, exhibit duplication of different anti-complement genes (i.e., duplication of c*spZ* and *bb_k32* in B379 and B408, respectively). Gene duplication has been shown to confer a selective advantage by increasing gene expression via the dosage effect (Kondrashov *et al*. 2002; Francino 2012). Our results thus raise an intriguing possibility that, while specific genes across *B. burgdorferi* strains may exhibit high sequence similarity and confer similar host-specific infection phenotypes (i.e., complement evasion), dosage effects imparted by variable gene copy numbers may help determine the overall host-specific phenotypes. In fact, Lyme borreliae genomes are known to harbor numerous gene families, suggesting that gene duplications are a common strategy for adapting to selective pressure and improving adaptive potential (Caine & Coburn 2016; Schwartz *et al*. 2021). A future study would investigate gene dosage effects and dosage compensation by deploying transcriptomics at different timepoints of infection. Further, our study added to the known diversity of methylation motifs involved in Restriction/Modification systems across *B. burgdorferi* strains that likely impact recombination across plasmids and/or differential gene expression (Rego *et al*. 2011; Casselli *et al*. 2018; Wachter *et al*. 2021). This supports the notion that the phenotypic variation among Lyme borreliae species or strains could also be driven by epigenetic differences in spirochetes. An examination of the expression differences among strains or the impact of horizontal gene transfer governed by Restriction/Modification systems would be essential for identifying epigenetic determinants of host association.

Our laboratory animal models and natural reservoir hosts are evolutionarily distant (house mice vs. white-footed mice diverged 24 Mya; quail and passerines, e.g. American robins, diverged 85 Mya (Steppan *et al*. 2004; Claramunt & Cracraft 2015)). This reflects the differences of spirochete-related phenotypes documented *in vitro* and *in vivo* between laboratory or reservoir hosts and the need to extend such a host association study to reservoir animals a future work (e.g. house mice vs. white-footed mice) (Barthold *et al*. 1991; Hanincová *et al*. 2008; Lin *et al*. 2022). Moreover, accurate assembly from short-read sequencing data of Lyme borreliae is challenging, as the genome of these species/strains include a large number of linear and circular plasmids, some of which are very conserved (cp32s) and/or constantly recombine (Casjens *et al*. 1997b, 2017). These characteristics highlight the importance of the multiple polygenic traits in impacting multiple phenotypes (e.g. host-specific complement evasion and dissemination), which requires a comparative genomics approach, despite several of single-and multi-locus typing schemes have been historically used to characterize genotypic variation (IGS, RST, MLST; Margos *et al*. 2008; Schwartz *et al*. 2021). Here we applied a multi-disciplinary approach to examine the concept of complement-mediated, host-specific infection, demonstrating the diversification of Lyme borreliae strains within single species. Comparing locus-specific phylogenetic vs. phenotypic differences, combined with comparative genomics, enables us to identify the potential determinants of host association. Such results can provide the evolutionary framing of the Lyme disease system, paving the road in dissecting the molecular mechanisms of host-pathogen interactions.

## MATERIALS AND METHODS

### Ethics statement

All mouse and quail experiments were performed in strict accordance with all provisions of the Animal Welfare Act, the Guide for the Care and Use of Laboratory Animals, and the PHS Policy on Humane Care and Use of Laboratory Animals. The protocol was approved by the Institutional Animal Care and Use Committee (IACUC) of Wadsworth Center, New York State Department of Health (protocol docket number 19-451. All efforts were made to minimize animal suffering.

### Mouse, quail, tick, bacterial strains, animal sera, and OmCI

BALB/c and Swiss Webster mice were purchased from Taconic (Hudson, NY). C3^-/-^ mice in BALB/c background were generated from the C3^-/-^(C57BL/6) purchased from Jackson Laboratory (Bar Harbor, ME) as described in our previous study (Hart *et al*. 2018). Common quail (*Coturnix coturnix*) were purchased from Cavendish Game Birds Farm (Springfield, VT). *Ixodes scapularis* tick larvae were purchased from the National Tick Research and Education Center, Oklahoma State University (Stillwater, OK) or from BEI Resources (Manassas, VA). Lyme borreliae-infected nymphs were generated as described in the section “Generation of infected ticks.” The *Borrelia* and *Escherichia coli* strains used in this study are described in Table 1 and S4. *E. coli* strains Rosetta Origami (DE3) (MilliporeSigma, Burlington, MA), and derivatives were grown in Luria-Bertani (BD Bioscience) broth or agar, supplemented with kanamycin (50µg/ml) or no antibiotics as appropriate. All *B. burgdorferi* strains were grown in BSK-II completed medium with no antibiotics and verified for their plasmid profiles for the strains that have the protocol established (Purser & Norris 2000) prior to the experiments and/or maintained in the passage less than 10 to avoid the confounding factors caused by plasmid missing. The sera from white footed mouse were obtained described previously (Lin *et al*. 2022), and quail sera were obtained from Canola Live Poultry Market (Brooklyn, NY). Prior to being used, these sera were screened with the C6 Lyme ELISA kit (Diamedix, Miami Lakes, FL) to determine whether the individual from which it was collected had prior exposure to *B. burgdorferi* by detecting antibodies against the C6 peptide of the *B. burgdorferi* protein VlsE (Lawrenz *et al*. 1999).

### Generation of recombinant OmCI and quail C8γ, and antisera against quail C8γ

To generate the plasmid in producing soluble OmCI proteins, a bi-cistronic expression plasmid was constructed to allow simultaneous production of the OmCI (NCBI Reference Sequence; AY560803.1; amino acids 19-168) and with mature human protein disulfide-isomerase (GenBank; X05130.1; amino acids 18-508) as described (Kuhn *et al*. 2016). The DNA fragment in this plasmid were first synthesized to include 1) the region 67-518 nucleotides encoding OmCI (*Ornithodoros moubata* complement inhibitor precursor), 2) a linker and a ribosome binding site (“GGAGGCAAAAA”), 3) a translational starting sites (“ATGAAA”), and 4) the mature human protein disulfide-isomerase from the five to three prime ends (Synbio Technology, Monmouth Junction, NJ). That DNA fragment was then added the restriction enzyme sites, BamHI and SalI, at 5’ and 3’ ends using PCR followed by being ligated into the vector, pET28a (MilliporeSigma), previously digested by respective restriction enzyme. The resulting plasmid was then transformed into the *E. coli* strain Rossetta Origami (DE3). To generate the plasmid in producing soluble γ chain of quail C8 (NCBI RefSeq XM_015878802.2; amino acid positions 50-228), the region 300-839 nucleotides encoding *c8g* (γ chain of C8 from common quail) were synthesized (Synbio Technology) and cloned in the same fashion. The resulting plasmid was also transformed into the *E. coli* strain Rossetta Origami (DE3). The histidine-tagged quail C8γ and OmCI were produced and purified by Ni-NTA affinity chromatography according to the manufacturer’s instructions (GE Healthcare, Piscataway, NJ). Antisera against quail C8γ were generated by immunizing four-week-old Swiss Webster mice with the recombinant quail C8γ as described (Benoit *et al*. 2011).

### Flow cytometry

The determination of mouse C5b-9 or quail C8 deposition using FACS was described previously with modifications (Hart *et al*. 2018; Marcinkiewicz *et al*. 2019a). In brief, PBS was used to wash spirochetes (1 x 10^8^ cells), which were then resuspended in the same buffer. The sera from mouse or quail were subsequently incubated with suspended spirochetes at a final concentration as 20% at 25°C for 1h. After incubation, spirochetes were washed by PBS and then resuspended in HBSC-DB (25mM Hepes acid, 150mM NaCl, 1mM MnCl_2_, 1mM MgCl_2_, 0.25mM CaCl_2_, 0.1% glucose, and 0.2% BSA). A rabbit anti-mouse C5b-9 polyclonal IgG (1:250x) (Complement Technology, Tyler, TX) and a mouse anti-quail C8 polyclonal serum (1:250x) generated in the section “Generation of recombinant quail C8γ, CspZ proteins, and antisera against quail C8γ” were used as primary antibody. An Alexa 647-conjugated goat anti-rabbit (ThermoFisher) or a goat anti-mouse IgG (ThermoFisher) (1:250x) was used as the secondary antibody. After staining, formalin (0.1%) was then added for fixing. The resulting fluorescence intensity of spirochetes was measured and analyzed by flow cytometry using a FACSCalibur (BD Bioscience) as described in the previous studies (Marcinkiewicz *et al*. 2019a).

### Serum resistance assays

The serum resistance of *B. burgdorferi* was measured as previously described, with modifications (Hart *et al*. 2018; Marcinkiewicz *et al*. 2019a; Lin *et al*. 2022). To determine the survivability of each of the *B. burgdorferi* strains in the sera, the mid-log phase of each of these strains were cultivated in triplicate and diluted to a final concentration of 5×10^6^ bacteria per milliliter into BSK-II medium without rabbit sera. The high passaged, non-infectious, and serum sensitive *B. burgdorferi* strain B313 was also included as control. The cell suspensions were mixed with sera collected from naïve white-footed mice or quail (60% (v/v) spirochetes and 40% (v/v) sera) in the presence or absence of 2µM of cobra venom factor (CVF) (Complement Technology) or recombinant OmCI. The sera pre-incubated at 65°C for 2h (heat-inactivated sera) were included as a control. To determine the impact of OmCI treatment to reduce the complement-mediated killing activity of quail, quail were subcutaneously injected with OmCI (1mg/kg of quail) or PBS buffer (control), and the sera were collected at 9 days post injection (dpi). *B. burgdorferi* strain B313 was cultivated and then mixed with those quail sera in the same fashion indicated above. The bacteria mixed with heat inactivated serum samples were also included as control. We determined the survivability of bacteria by counting the number of motile and immotile bacteria, in which we have shown to yield similar results to the methodology counting the bacteria based on live: dead staining (Marcinkiewicz *et al*. 2019b; Frye *et al*. 2020) Basically, the number of motile spirochetes was measured under dark field microscopy at 0 and 4h following incubation with sera. The percent survival of *B. burgdorferi* was calculated by the normalization of motile spirochetes at 4h post incubation to that immediately after incubation with sera.

### Evolutionary genomics

To evaluate the correlation between phylogenetic signal and serum survivability phenotypes, we compiled genomic sequences for each of the 20 *B. burgdorferi* strains as well as for the *Borrelia bissettiae* strain DN127 (Table 1). Of these, fully or partially assembled genomes of 11 strains were available on Genbank: JD1, MM1, CA1-11.2A, ZS7, PAli, PAbe, B31, cN40, B331, WI91-23, and 29805 (Accession numbers in Table 1). One strain, PMeh, was unassembled but represented by raw Illumina reads on the Sequence Read Archive (SRA) (Table 1). Genomic DNA for four strains, B356, N40-D10/E9, 297, and Bbss62, was extracted from cultures. Library preparation was carried out with the Nextera DNA Flex Library Preparation kit [Illumina, San Diego, CA] and sequenced on a MiSeq instrument using the v2 500 cycle kit at the Advance Genomic Technologies Cluster, Wadsworth Center (Albany, NY). Raw reads for PMeh, B356, N40-D10/E9, 297, and Bbss62 were quality-filtered and trimmed using the *trim_galore* wrapper script specifying a minimum quality score of 20 and a minimum length of 60 nucleotides (https://www.bioinformatics.babraham.ac.uk/projects/trim_galore). Assembly took place in SPAdes v3.15.3 using the --isolate mode (Bankevich *et al*. 2012). Sequencing and assembly details for B31-5A4, B408, and B379 are described below.

The linear chromosome of *B. burgdorferi* contains genes essential for biological functions and are subjected to strong purifying selection, providing a useful representation of neutral population genetic relationships among strains (Tyler *et al*. 2018). We extracted core genes found across each of the 20 *B. burgdorferi* genomes and the *B. bissettii* strain DN127 using Roary v3.13.0 (parameters: -i 0.9, -e, -s), after annotation with PROKKA v1.14.6 (Seemann 2014). We used IQTree v1.6.12 to estimate a phylogenetic tree from the core set of chromosomal genes, partitioning by gene, implementing a full substitution model testing procedure, along with the ultrafast bootstrap procedure and the Shimodaira-Hasegawa-like approximate likelihood ratio test to gauge internode branch support, using the following parameters: --bb 10000 -alrt 10000 -safe - p -t RANDOM (Nguyen *et al*. 2015).

The phylogenetic history of several genes known to be involved in host complement evasion by *B. burgdorferi* including *ospC*, *cspZ*, *cspA*, and *bb_k32* was reconstructed. We noted that the strains 29805, N40-D10/E9, and B356 are missing *bb_k32* alleles and thus did not include those alleles in the *bb_k32*-derived phylogeny. Sequences for each of these genes were aligned as translated amino acids using MAFFT v7.480 (Katoh & Standley 2013) and phylogenies were constructed using IQTree v1.6.12 and the same parameters listed above. We calculated Pagel’s lambda, a measure of phylogenetic signal (Molina-Venegas & Rodríguez 2017), specifying anti-complement phenotype as a discrete trait, using the *geiger* R package (Pennell *et al*. 2014). To evaluate analytical consistency, we ran each analysis 100 times and built a distribution of resulting λ values.

### Generation of infected ticks

Generating infected *I. scapularis* ticks has been described previously with modifications (Kern *et al*. 2016). BALB/c C3-deficient mice were infected intradermally with 10^5^ of each of the *B. burgdorferi* strains. The DNA extracted from the ear tissues was applied to qPCR to verify DNA positivity of *B. burgdorferi* as verification of infection (see section “Quantification of spirochete burden”). The uninfected *Ixodes scapularis* larvae (∼100 to 200 larvae per mouse) were allowed to feed to repletion on the infected mice as described previously (Hart *et al*. 2021). The engorged larvae were collected and permitted to molt into nymphs in a desiccator at room temperature with 95% relative humidity and light dark control (light to dark, 16:8 hours).

### Intravenous inoculation of *B. burgdorferi*

The short-term intravenous inoculation experiments were performed, as described with modifications (Caine & Coburn 2015; Caine *et al*. 2017). Four- to six-week-old male and female BALB/c or C3^-/-^ mice in BALB/c background were inoculated with 1×10^7^ *B. burgdorferi* cells in 100 µl via lateral tail veins, whereas four- to six-week-old male and female untreated or OmCI-quail were injected with 1×10^8^ *B. burgdorferi* cells in 100 µl via brachial veins in the wing. At one hour after inoculation, mice were euthanized to collect blood by cardiac puncture while the blood from quail was collected from the brachial vein from the other side of wing. DNA isolated from the blood samples was used for quantitation of *Borrelia* by qPCR described below (Caine & Coburn 2015; Caine *et al*. 2017).

### Mouse and quail infection by ticks

The flat nymphs were placed in a chamber on four- to six-week old male and female BALB/c or C3^-/-^ mice in BALB/c background or four- to six-week old male and female untreated or OmCI-quail as described previously (Hart *et al*. 2021). For OmCI-treated quail, the quail were subcutaneously injected with OmCI (1mg/kg of quail) a day prior to the nymph feeding. The engorged nymphs were obtained from the chambers at four days post tick feeding. Blood, tick placement site of skin, and heart from the quail or mice, bladder and tibiotarsus joints from the mice, and brain from the quail were collected at 9- (for quail), 10-, or 14- (for mice) days post nymph feeding.

### Quantification of spirochete burden

The DNA from tissues, blood, or ticks was extracted as described previously (Hart *et al*. 2021). qPCR was then performed to quantitate bacterial loads. Spirochete genomic equivalents were calculated using an ABI 7500 Real-Time PCR System (ThermoFisher Scientific) in conjunction with PowerUp SYBR Green Master Mix (ThermoFisher Scientific) based on amplification of the Lyme borreliae 16S rRNA gene using primers BB16srRNAfp and BB16srRNArp (Table S5) with the amplification cycle as described previously (Marcinkiewicz *et al*. 2019a). The number of *recA* copies was calculated by establishing a threshold cycle (Cq) standard curve of a known number of *recA* gene extracted from *B. burgdorferi* strain B31-5A4, then comparing the Cq values of the experimental samples.

### Comparative genomics

<u>1) Sequencing and assembly:</u> B31-5A4, B379, and B408 were grown for genomic sequencing as described above, and genomic DNA was extracted using a cetyltrimethylammonium bromide and organic extraction method (Xu *et al*. 2015). The genomic DNA was assessed for shearing with gel electrophoresis. Genomic libraries were independently prepared for sequencing using the SMRTbell library prep kit, with a mean sheared genomic DNA distribution in the range 4-6kb, then sequenced on a PacBio RSII system at the Icahn School of Medicine at Mount Sinai Genomics Core Facility (New York, NY). Genomes were assembled using the default HGAP3 pipeline. Upon inspection, all three assembled genomes revealed evidence of “adapter skipping” artifacts in the form of long (5-20kb) inverted repeats found only at the termini of many linear plasmids, a known issue commonly identified across *Borrelia* genomes (Schwartz *et al*. 2021). Adapter skipping occurs when one hairpin adapter fails to ligate during library preparation, causing individual subreads to contain both sense and antisense sequences (pers, commun. with PacBio Field Applications Scientist). While the *recalladapters* script is now available to correct this issue, it is not compatible with RSII system-derived sequences. We therefore regrew the cultures and prepared genomic DNA for each strain as described above and sequenced each on a Sequel I instrument. Initial assembly with HGAP4 using default parameters revealed nearly identical adapter skipping artifacts, which were only ameliorated after implementing custom subread filtering scripts provided by PacBio’s software development team (Text S1).

High Fidelity (HiFi) reads were next generated subread consensus sequence for each strain using Pacbio’s *ccs* script (github.com/PacificBiosciences/pbbioconda). Genomes for each strain were assembled using Canu v2.1 using the following parameters: -pacbio-hifi, minReadLength=1500, minOverlapLength=1500 (Nurk *et al*. 2020). Circular contigs identified by Canu were trimmed and rotated to match the start position of reference sequences B31, or of JD1 and cN40 for plasmids not found in B31, using Gepard v1.40 and the *fasta_shift* script available in the fasta_tools package (github.com/b-brankovics/fasta_tools). Plasmids were identified by homology comparisons to known PFam32 loci, plasmid partitioning genes commonly used to identify *Borrelia* plasmids (Casjens *et al*. 2012), and based on Gepard v1.40 dotplot and BLASTn matches to known plasmids from B31, JD1 and N40 reference genomes. Genomes were annotated with PROKKA v1.14.6 (Seemann 2014) using proteins from the B31 reference genome for initial comparison (Accession: GCA_000008685.2).

<u>2) Base modifications:</u> For each genome, base modifications and motifs were detected using Pacific Bioscience’s SMRT analysis pipeline, where modification calls were only retained if they had a quality value of 400 or greater for B31-5A4 and B379, or 200 or greater for B408. Quality value thresholds were determined from breaks observed in the quality value distributions, as suggested by PacBio Product Application Specialists (Fig. S5). Positions of methylated motifs were extracted and a 1000bp sliding window was used to visualize and compare genome wide methylation rates using the *zoo* R package (Zeileis & Grothendieck 2005).

<u>3) Plasmid and gene comparison:</u> Shared plasmids between B31-5A4, B379, and B408 were compared using BLAST sequence alignments, gene annotations, and the number of methylated m6A motifs per 1000bp with Easyfig (Sullivan *et al*. 2011). To identify nucleotide similarity among annotated genes, we used *NUCmer* and *mummerplot* (using parameter -c for percent identity plots) from the MUMmer v3.1 package (Kurtz *et al*. 2004).

### Statistical analysis

Significant differences between samples were assessed using the Mann-Whitney *U* test or the Kruskal-Wallis test with the two-stage step-up method of Benjamini, Krieger, and Yekutieli. A P-value < 0.05 (*) or (^#^) was considered to be significant (Benjamini *et al*. 2006).

## ACKNOWLEDGEMENTS

We thank Patricia Rosa, John Leong, George Chaconas, Richard Marconi, Robert Gilmore, Nikhat Parveen, Peter Kraiczy, Melissa Caimano, and Nicholas Mantis for providing *B. burgdorferi* and *E. coli* strains, and Sanjay Ram for providing the mouse strain. We also thank Simon Starkey, Dierdre Torrisi and other staff in Wadsworth Animal Core for assistance with Animal Care, ATGC core for sanger sequencing and illumina short read sequencing, Renjie Song at Wadsworth Biochemistry and Immunology Core for flow cytometry, Wadsworth Center Media & Tissue Culture Core for preparation of *E. coli* and Lyme borreliae culture media, and the Genomics Core Facility at Icahn School of Medicine at Mount Sinai for PacBio long read sequencing. This work was supported by NSF IOS1755370 (MC and MDW), NSF IOS1754995 (SOK), NSF IOS1755286 (ALM, ADII, PL, TAN, JLS, ATC, and YPL) and New York State Department of Health Wadsworth Center Start-Up Grant (KS, ALM, PL, TAN, and YPL). The funders had no role in study design, data collection, interpretation, or the decision to submit the work for publication. The authors have no conflict of interest to declare.

