## Supplementary information for "Phylogenomic diversity elucidates mechanistic insights into Lyme borreliae host association"

**SUPPLEMENTAL FIGURE LIGENDS**

**Figure S1. Distribution of Pagel's lambda values for 100 iterations for the correlation of the anti-complement gene phylogeny and the phenotypes of host-specific complement evasion.** Histograms indicates the certainty of Pagel's lambda estimation for the analysis of host-specific anti-complement phenotype from Figure 1 paired with the phylogeny of **(A)** *cspA*, **(B)** *bb_k32*, **(C)** *cspZ*, **(D)** *ospC*, and **(E)** chromosome core genes. Only estimations for *ospC* and the core chromosomal genes result in stable estimations.

**Figure S2. The tested *B. burgdorferi* strains in this study exhibited similar burdens in flat and fed nymphs.** C3-deficient BALB/c ($C3^{-/-}$ BALB/c) mice were intradermally inoculated at the dose of $10^5$ spirochetes per milliliter of BSK media without rabbit sera with *B. burgdorferi* strains B31-5A4, B379, or B408. At 14 days post infection, the uninfected *I. scapularis* larval ticks were allowed to feed on each of these mice until they are replete. After the replete larvae molt into flat nymphs, those *B. burgdorferi*-infected flat nymphs were allowed to feed on **(B)** BALB/c mice, **(C)** untreated quail, **(D)** C3-/- BALB/c mice, or **(E)** OmCI-treated quail (OmCI-quail) to repletion. The bacterial loads in **(A)** flat nymphs, or **(B to E)** replate nymphs were determined by qPCR. Shown are the geometric mean ± geometric standard deviation of eight flat nymphs or nymphs feeding on BALB/c mice, seven nymphs feeding on quail or $C3^{-/-}$ BALB/c mice, or ten nymphs feeding on OmCI-treated quail (except for B408-infected nymphs feeding on OmCI-treated quail, there were eight nymphs per group). There was no statistical difference ($p > 0.05$) of the spirochete burdens among different groups of the replete ticks using Kruskal-Wallis test with the two-stage step-up method of Benjamini, Krieger, and Yekutieli.

**Figure S3. Complement is not sufficient to dictate *B. burgdorferi* genotype-specific bloodstream survival and tissue colonization in mice at 14dpf.** The *I. scapularis* nymphs carrying *B. burgdorferi* strains B31-5A4, B379, or B408, were allowed to feed until they are replete on five **(A to E)** BALB/c mice or **(F to J)** C3$^{-/-}$ mice in BALB/c background. The mice were euthanized at 14-days post nymph feeding (dpf), respectively. The bacterial loads at **(A, F)** the site where nymphs fed (“inoc. site”), **(B, G)** blood, **(C, H)** tibiotarsus joints, **(D, I)** bladder, and **(E, J)** heart of mice collected immediately after euthanasia were determined by qPCR. The bacterial loads in tissues or blood were normalized to 100 ng total DNA. Shown are the geometric mean of bacterial loads ± geometric standard deviation of five mice or quail per group. Significant differences ($p < 0.05$, the Kruskal-Wallis test followed by the two-stage step-up method of Benjamini, Krieger and Yekutieli.) in the spirochete burdens between two strains relative to each other (‘‘#’’).

**Figure S4. No significant sequence variation, deletion, or duplication are found in cp26 among B31-5A4, B379, and B408.** The sequence of each plasmid is represented by a black line and gene annotations are depicted by teal arrows, labeled according to homology to known genes in *B. burgdorferi* strain B31. Segments connecting each strain represent filtered BLAST results in either the same orientation (blue) or opposite orientation (red), with darker shades representing closer matches. Orange graphs above each strain depict the sliding window calculation of methylated nucleotides per 1000 bases. The red arrows indicated the loci of *ospC*.

**Figure S5. Base modification detection diagnostic plots for quality score cutoff.** Distributions describe the confidence in the observed base modification in relation to overall sequencing depth for *B. burgdorferi* strains **(A)** B31-5A4, **(B)** B379, and **(C)** B408. Natural breaks in the distribution provide insight into appropriate QV cutoff scores for each strain.

### SUPPLEMENTARY TABLES

**Table S1. Long-read sequencing measures of *B. burgdorferi* strains B31-5A4, B379, and B408.**

| | B31-5A4 | B379 | B408 |
|---|---|---|---|
| **Total base pairs sequenced** | 722,553,355 | 357,749,072 | 568,303,383 |
| **Number of reads** | 72,321 | 49,524 | 78,429 |
| **N50 read length (bp)** | 13,854 | 10,132 | 10,078 |
| **Mean read length (bp)** | 9,990 | 7,223 | 7,246 |
| **Total length of contigs (bp)** | 1513692 | 1417697 | 1392053 |
| **Total length of plasmids (bp)** | 604037 | 496924 | 488804 |
| **Number of total contigs** | 20 | 18 | 18 |
| **Number of chromosomal contigs** | 1 | 1 | 1 |
| **Number of linear plasmids** | 11 | 8 | 10 |
| **Number of circular plasmids** | 9 | 9 | 7 |
| **Number of annotations** | 1527 | 1435 | 1408 |
| **Methylations (per 1000 bp)** | 6.11 | 4.63 | 2.78 |

**Table S2. The methylation motifs identified in B31-5A4, B379, and B408.**

| Strain | Motif Sequence | Center Position | Modification Type | Proportion of Motifs Methylated | Number of Motifs Methylated | Number of Motifs in Genome |
|---|---|---|---|---|---|---|
| **B31-5A4** | CGRKA | 5 | m6A | 0.948 | 2088 | 2202 |
| **B379** | GNAAGC | 4 | m6A | 0.982 | 1871 | 1906 |
| **B408** | GAAGG | 3 | m6A | 0.967 | 1434 | 1483 |

**Table S3. Variable genes from locus-specific comparison in B31-5A4, B379, and B408.**

| Nucleotide identity (%) | B408 vs. B31-5A4 | | | |
|---|---|---|---|---|
| | B408 | | B31-5A4 | |
| 80.25 | BB_G29 | Hypothetical protein | BB_G29 | Hypothetical protein |
| 82.21 | BB_M38 | ErpK protein | BB_M38 | ErpK protein |
| 82.24 | BB_I01 | Hypothetical protein | BB_0408 | Hypothetical protein |
| 82.36 | BB_Q35 | Mlp family lipoprotein | BB_Q35 | Mlp family lipoprotein |
| 82.41 | BB_H09 | Class I SAM-dependent DNA methyltransferase | BB_H09 | Class I SAM-dependent DNA methyltransferase |
| 82.84 | BB_M29 | ERF family protein | BB_O29 | DUF188 domain-containing protein |
| 82.87 | BB_B23 | NCS2 family permease | BB_B22 | NCS2 family permease |
| 83.2 | BB_E02 | Class I SAM-dependent DNA methyltransferase | BB_H09 | Class I SAM-dependent DNA methyltransferase |
| 83.42 | BB_C11 | Site-specific integrase | BB_Q45 | Site-specific integrase |
| 83.5 | BB_E02 | Class I SAM-dependent DNA methyltransferase | BB_E02 | Class I SAM-dependent DNA methyltransferase |

| | | | | |
|---|---|---|---|---|
| 83.75 | BB_Q35 | Mlp family lipoprotein | BB_Q35 | Mlp family lipoprotein |
| 84.05 | BB_H09 | Class I SAM-dependent DNA methyltransferase | BB_E02 | Class I SAM-dependent DNA methyltransferase |
| 84.48 | BB_E02 | Class I SAM-dependent DNA methyltransferase | BB_H09 | Class I SAM-dependent DNA methyltransferase |
| 84.97 | BB_H09 | Class I SAM-dependent DNA methyltransferase | BB_E02 | Class I SAM-dependent DNA methyltransferase |

| **Percent identity** | **B379 vs. B31-5A4** | | | |
|---|---|---|---|---|
| | **B379** | | **B31-5A4** | |
| 80.2 | BB_K13 | SIMPL domain-containing protein | BB_K13 | SIMPL domain-containing protein |
| 80.25 | BB_G29 | Hypothetical protein | BB_G29 | Hypothetical protein |
| 81.73 | BB_C03 | Chromosome replication/partitioning protein | BB_L34 | Chromosome replication/partitioning protein |
| 82.24 | BB_I01 | Hypothetical protein | BB_0454 | Hypothetical protein |
| 82.43 | BB_B22 | NCS2 family permease | BB_B23 | NCS2 family permease |
| 82.53 | BB_B23 | NCS2 family permease | BB_B22 | NCS2 family permease |

| | | | | |
|---|---|---|---|---|
| 82.63 | BB_Q35 | Mlp family lipoprotein | BB_Q35 | Mlp family lipoprotein |
| 83.41 | BB_Q35 | Mlp family lipoprotein | BB_Q35 | Mlp family lipoprotein |
| 83.7 | BB_L27 | Hypothetical protein | BB_Q34 | Hypothetical protein |
| 84.1 | BB_J34 | Lipoprotein | BB_J34 | Lipoprotein |
| 84.27 | BB_E02 | Class I SAM-dependent DNA methyltransferase | BB_H09 | Class I SAM-dependent DNA methyltransferase |
| 84.73 | BB_L27 | Hypothetical protein | BB_Q34 | Hypothetical protein |

| **Percent identity** | **B379 vs. B408** | | | |
|---|---|---|---|---|
| | **B379** | | **B408** | |
| 79.71 | BB_K13 | SIMPL domain-containing protein | BB_K13 | SIMPL domain-containing protein |
| 80.38 | BB_I16 | Virulence associated lipoprotein | BB_0397 | Hypothetical Protein |
| 81.62 | BB_E02 | Class I SAM-dependent DNA methyltransferase | BB_H09 | Class I SAM-dependent DNA methyltransferase |
| 81.84 | BB_Q24 | DUF693 family protein | BB_O17 | DUF693 family protein |
| 81.84 | BB_Q24 | DUF693 family protein | BB_O17 | DUF693 family protein |
| 81.84 | BB_Q24 | DUF693 family protein | BB_O17 | DUF693 family protein |

| | | | | |
|---|---|---|---|---|
| 82.44 | BB_H09 | Class I SAM-dependent DNA methyltransferase | BB_H09 | Class I SAM-dependent DNA methyltransferase |
| 82.53 | BB_N28 | Mlp family lipoprotein | BB_N28 | Mlp family lipoprotein |
| 82.62 | BB_B23 | NCS2 family permease | BB_B22 | NCS2 family permease |
| 82.65 | BB_E02 | Class I SAM-dependent DNA methyltransferase | BB_E02 | Class I SAM-dependent DNA methyltransferase |
| 82.76 | BB_H09 | Class I SAM-dependent DNA methyltransferase | BB_E02 | Class I SAM-dependent DNA methyltransferase |
| 82.87 | BB_B22 | NCS2 family permease | BB_B23 | NCS2 family permease |
| 82.95 | BB_Q24 | DUF693 family protein | BB_O17 | DUF693 family protein |
| 82.95 | BB_Q24 | DUF693 family protein | BB_O17 | DUF693 family protein |
| 83.1 | BB_Q35 | Mlp family lipoprotein | BB_Q35 | Mlp family lipoprotein |
| 83.13 | BB_A24 | Decorin-binding protein DbpA | BB_A24 | Decorin-binding protein DbpA |
| 83.19 | BB_M34 | Hypothetical protein | BB_M34 | Hypothetical protein |
| 84.05 | BB_E02 | Class I SAM-dependent DNA methyltransferase | BB_H09 | Class I SAM-dependent DNA methyltransferase |

| | | | | |
|---|---|---|---|---|
| 84.2 | BB_J34 | Lipoprotein | BB_J34 | Lipoprotein |
| 84.71 | BB_N42 | DUF603 domain-containing protein | BB_S44 | DUF603 domain-containing protein |

**Table S4. The *E. coli* strains and the plasmid used in this study**

| Strain or plasmid | Genotype or characteristic | Source |
|---|---|---|
| *E. coli* | | |
| DH5α | F- Φ80lacZΔM15 Δ(lacZYA-argF) U169 recA1 endA1 hsdR17(rk-, mk+) phoA supE44 thi-1 gyrA96 relA1 λ- | ThermoFisher |
| Rosetta-gami(DE3) | F– ompT hsdSB (rB– mB–) gal dcm lacY1 ahpC (DE3) gor522::Tn10 trxB pRARE (CamR, KanR, TetR) | MilliporeSigma |
| Rosetta-gami(DE3)/pET28a-qC8γ-hPDI | Rosetta-gami(DE3) producing histidine-tagged residues 50 to 228 of the γ chain of quail C8 and residues 18 to 508 of human protein disulfide isomerase. | This study |
| Rosetta-gami(DE3)/pET28a-OmCI-hPDI | Rosetta-gami(DE3) producing histidine-tagged residues 19 to 168 of OmCI and residues 18 to 508 of human protein disulfide isomerase. | This study |
| Plasmids | | |
| pET28a | KanR[a]; Histidine-tagged protein expression vector | EMD Milliopore |
| pET28a-qC8γ-hPDI | KanR; pET28a encoding histidine protein residues 50 to 228 of the γ chain of quail | This study |

| | | |
|---|---|---|
| | C8 with residues 18 to 508 of human protein disulfide isomerase. | |
| pET28a-OmCI-hPDI | KanR; pET28a encoding histidine protein residues 19 to 168 of OmCI with residues 18 to 508 of human protein disulfide isomerase. | (Kuhn *et al.* 2016) |

[a] Kanamycin resistant

**Table S5. Primers used in this study.**

| Primer | Sequence[a] | Amplified DNA fragment | Source |
|---|---|---|---|
| BBCspZ_prt_fp | GCGGATCCGATGTTAGTAGATTAAATC | *cspZ* without the signal peptide from *Borrelia burgdorferi* | (Marcinkiewicz *et al.* 2019b) |
| BBCspZ_prt_rp | GCGTCGACCTATAATAAAGTTTGCTTA | | |
| BBK32_full_fp | TGGAATCCGACTTAAAATGATTTAA | *bbk32* from *Borrelia burgdorferi* | This study |
| BBK32_full_rp | CATATTTACATATTATGTAGCCTG | | |

[a] Restriction enzyme sites are underlined

### SUPPLEMENTARY TEXT

**Text S1 Scripts used for additional subread filtering to remove adapter skipping reads from subread dataset, provided by Pacific Biosciences product application specialists and developer team.**

```
samtools view BC19.subreads.bam \
| awk '{split($23,a,":"); if(and(a[3],64)||and(a[3],128)){print $1}}' \
| tr '/' ' ' | awk '{print $2}' | sort -n | uniq > bad_adapters_holes.txt
bamsieve --blacklist bad_adapters_holes.txt BC19.subreads.bam BC19.filtered.subreads.bam
ccs --log-level INFO -j 16 BC19.filtered.subreads.bam BC19.filtered.ccs.bam
bam2fastq -o BC19.filtered.ccs BC19.filtered.ccs.bam
hifiasm -o BC19.asm -t16 -f0 BC19.filtered.ccs.fastq.g
```
