## Supplementary figures and images for "Phylogenomic diversity elucidates mechanistic insights into Lyme borreliae host association"

### Fig. S1

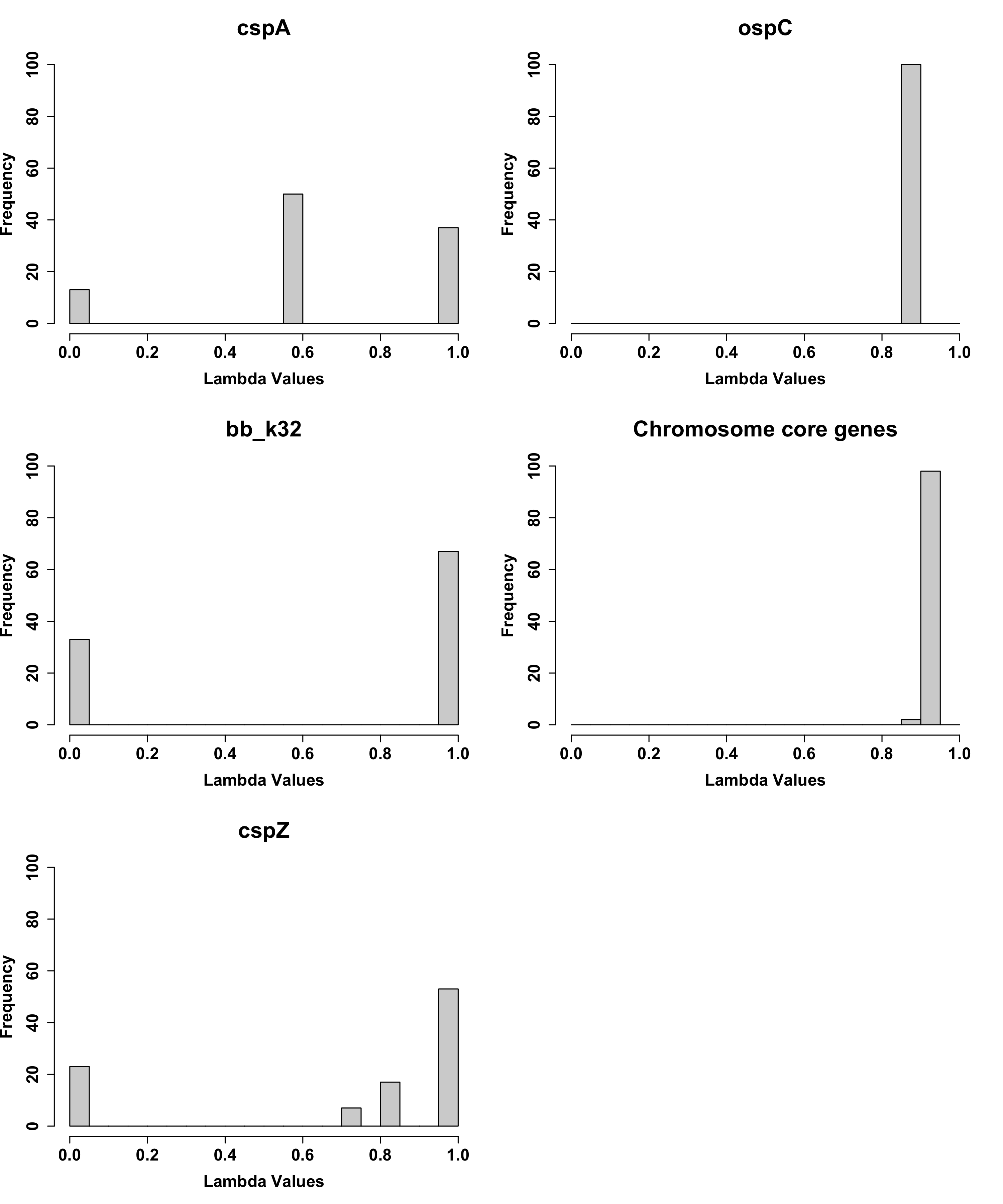

### Fig. S2

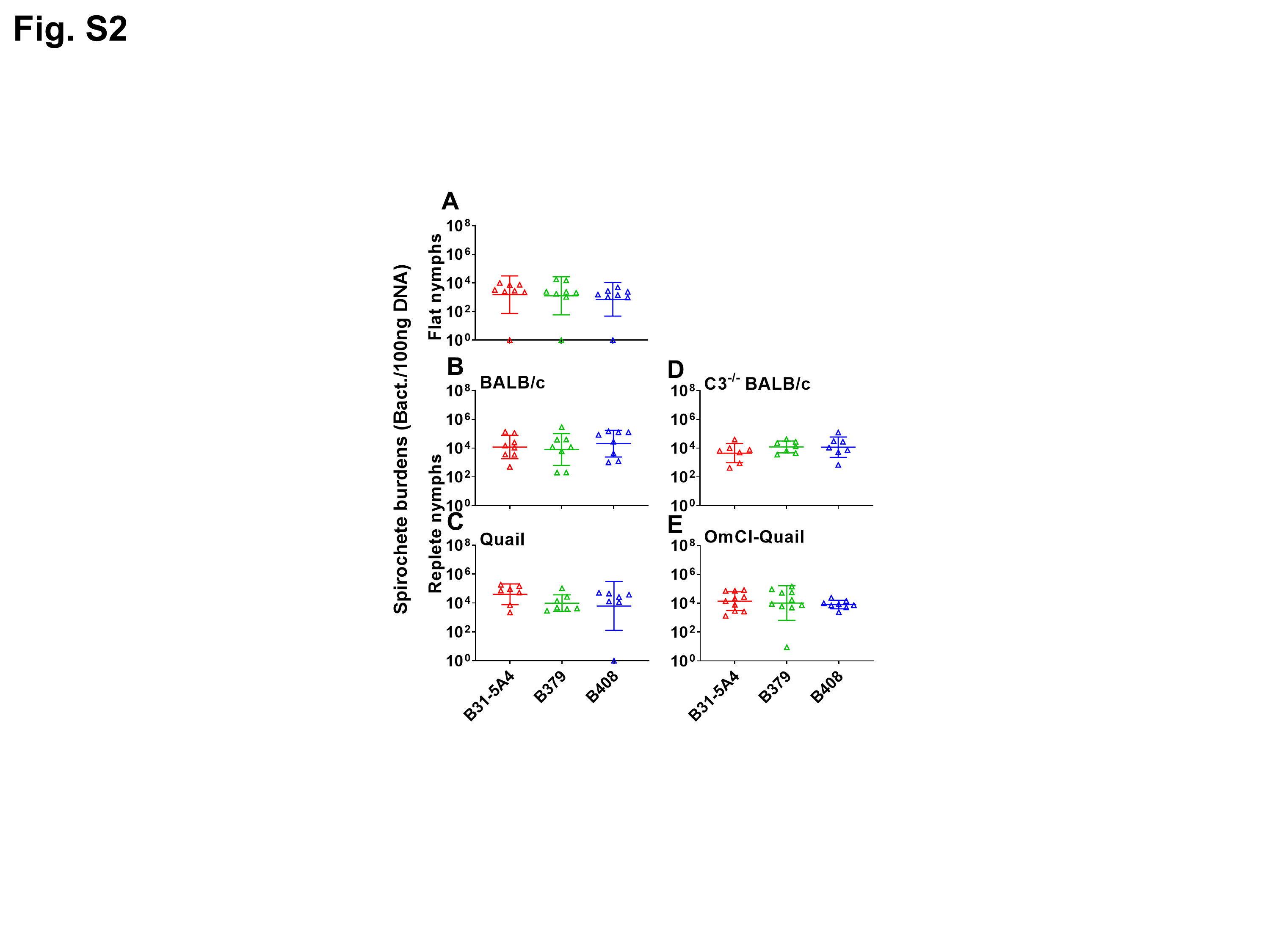

### Fig. S3

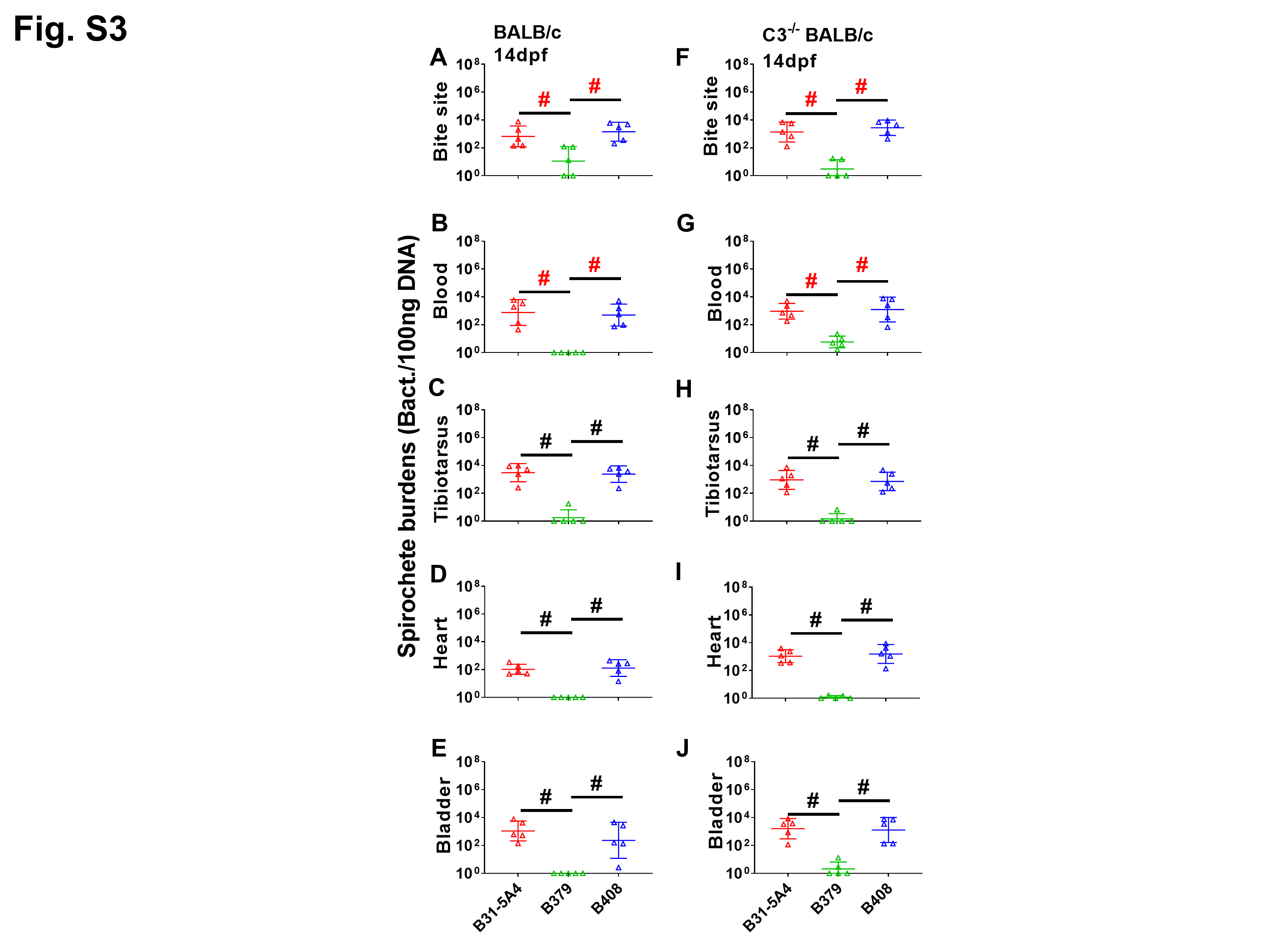

### Fig. S4

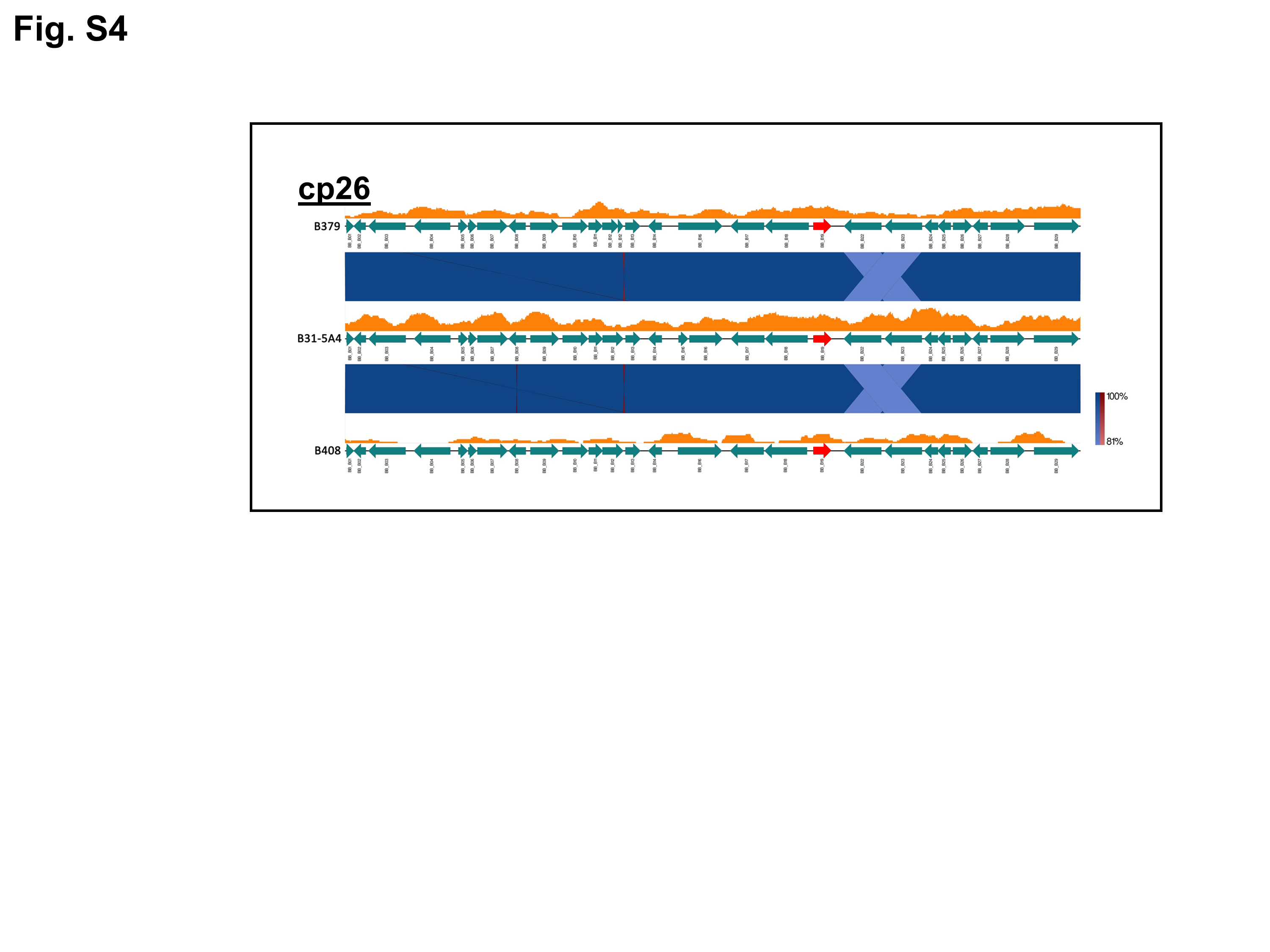

### Fig. S5

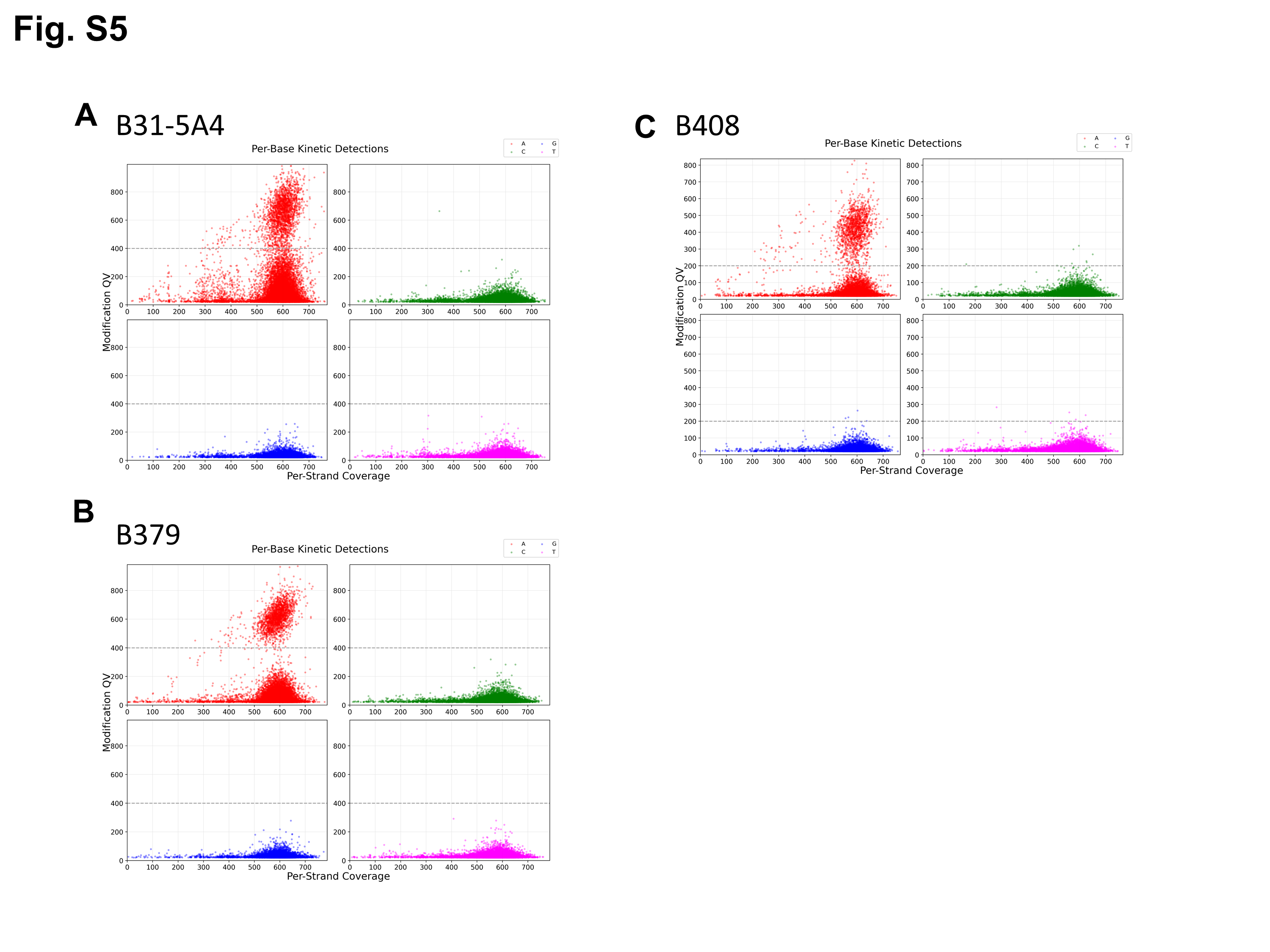
